# Integrating functional and phylogenetic perspectives reveals new priorities for bird and mammal conservation

**DOI:** 10.64898/2025.12.10.690976

**Authors:** Lucy Somekh, John N Griffin, Catalina Pimiento, William D Pearse

## Abstract

Finite resources dictate that conservation biologists must prioritise some species over others. Conservation metrics, such as EDGE (Evolutionarily Distinct and Globally Endangered; EDGE1) and FUSE (Functionally Unique, Specialised and Endangered), make prioritisation based on species’ evolutionary or functional distinctiveness, respectively, and degree of threat. EDGE is in part based on the rationale that the evolutionary distinctiveness it captures serves to maximise biodiversity in form and function. However, doubts exist as to whether or not evolutionary distinctiveness truly serves as a proxy for functional distinctiveness, and, therefore, if the prominent use of EDGE lists by conservation practitioners adequately protects functional distinctiveness. To address this, we conducted a direct global comparison of EDGE against FUSE. Unlike EDGE, FUSE directly measures and prioritises functional distinctiveness. Here, we compare EDGE and FUSE scores for two well-studied groups: mammals (*n* = 5319 species) and birds (*n* = 7932 species). These groups are central to global conservation and, until now, have not been assessed under FUSE. We find that species rankings under EDGE and FUSE differ significantly, highlighting that these two metrics capture distinct, complementary aspects of biodiversity. This suggests that conservation strategies based solely on EDGE may overlook species with critical functional roles. Rather than assuming alignment between evolutionary and functional distinctiveness, we propose integrating both within a single measure, leveraging the strengths and benefits of each. To this end, we present a new conservation metric – EFUSE (Evolutionarily and Functionally Unique, Specialised, and Endangered) – which incorporates both evolutionary and functional distinctiveness into a single measure. EFUSE ensures that important components of biodiversity, which relate to ecosystem functioning, nature’s future contributions to people, and the intrinsic value of species, are adequately maximised in conservation decision-making.

## Introduction

Global biodiversity is currently declining at a rate several orders of magnitude higher than background estimates from the fossil record (1–3). This scale of decline, combined with finite resources, is forcing conservationists to make difficult decisions over which species should be the primary focus of conservation efforts (4, 5). To aid this decision-making, numerous methods of species prioritisation have been proposed (e.g. 6, 7-9). These methods largely aim to make the decision-making over which species or geographical areas should form the focus of conservation resources a more objective process. This would ensure that human bias towards, for example, charismatic species, does not prevent the conservation of the most important and valuable species. However, defining important and valuable species is inherently subjective, making the process of selecting which prioritisation method to utilise subjective also (10). This is exemplified by a subset of these methods – conservation metrics – which are used to construct lists of species and/or geographical areas with high conservation value, based on attributes such as threat level, distinctiveness, or feasibility of conservation. Species-based metrics largely assign high priority to species whose existence maximises biological diversity in form and function, in order to keep humanity’s options open by preserving a species assemblage which is as diverse as possible (4, 11–13). However, conservation metrics measure species’ contributions to biodiversity in different ways and can, as such, largely be split into two categories: those which measure evolutionary diversity and those which measure functional (trait) diversity. An understanding of how these metrics correlate with or substitute each other will inform their utility for prioritizing conservation (4). Should these metrics correlate poorly, the use of one over the other may result in important species being insufficiently protected.

The only evolutionary-based conservation metric with a dedicated conservation program is EDGE (7, 14). The EDGE of Existence has funded at least 97 projects in 46 countries to date (15, 16). EDGE has also been adopted as a complementary indicator for monitoring the progress within the Kunming-Montreal Global Biodiversity Framework, which has set the 30 × 30 target, aiming to have at least 30% of the world’s land and sea areas effectively conserved and managed by 2030 (COP15 United Nations Biodiversity Conference 2022). The original EDGE protocol (7) has since been updated to EDGE2 (17). However, both approaches combine a measure of evolutionary distinctiveness (ED) with global endangerment (GE). In both versions, ED is measured in terms of phylogenetic diversity (PD; 18), calculated by partitioning the phylogenetic branch lengths spanning a set of taxa. The popularity of the EDGE metric is, in part, based on the fact that phylogenies (needed to calculate PD) are readily available for a wide variety of taxa (19). By comparison, the availability of direct measures of FD, such as trait data, is relatively limited (20). Proponents of EDGE believe that if evolutionary history captures variation in form and function, then preserving maximum PD will also serve to maximise functional diversity (FD), i.e., biological diversity in form and function – this has been termed the ‘phylogenetic gambit’ (20, 21). More recently, however, as trait data has become more readily available, some conservationists have proposed metrics which measure and aim to maximise functional trait diversity directly (9, 22). The FUSE metric, for example, incorporates the functional uniqueness, functional specialisation and GE of species (9). In measuring multiple dimensions of species’ function, FUSE allows us to generalise species’ functional contributions to ecosystems and contemplate the potential ecological consequences of their extinction.

Understanding the congruence between evolutionary- and functional-based metrics (like EDGE and FUSE, respectively) is essential to developing an understanding of how subjective decisions in conservation impact how and where finite resources are directed. EDGE is being used in-part based on the belief that ED serves as a proxy for PD, but previous research into the phylogenetic gambit raises questions about this belief and therefore the alignment between EDGE and FUSE scores. If ED were a reliable proxy for FD, we would expect a high degree of congruence between EDGE and FUSE. However, evidence suggests that alignment between ED and FD is often weak or inconsistent, varying across clades and geographical contexts (20, 21). Such variation likely reflects differences in evolutionary histories (23, 24).

For instance, adaptive radiations can result in species that have quickly diverged and are thus functionally dissimilar, despite their phylogenetic closeness (25), whereas convergent evolution can yield functionally similar species that are phylogenetically distant (26). In contrast, ED and FD tend to align more closely in instances where trait evolution follows a strong phylogenetic signal (27). Collectively, these findings underscore that ED serves as a useful surrogate for FD only when functional traits reliably mirror evolutionary history.

To truly understand the congruencies between EDGE and FUSE lists, and the impacts incongruencies will have on conservation, these lists must be calculated for a wider range of taxa and directly compared. To date, a direct, empirical comparison of EDGE and FUSE scores has only been conducted for elasmobranchs (sharks, rays and skates), with limited congruence observed and just two species in common among the top-20 lists (28). In this paper, we examine the relationship between EDGE and FUSE using all extant mammals and birds, for which evolutionary and functional (trait) data are readily available. Our analysis reveals limited congruence between the two metrics, with significant potential implications for conservation prioritisation. Having identified this disparity and after evaluating the respective strengths and limitations of EDGE and FUSE, we developed a new metric which integrates features of them both. We propose EFUSE (Evolutionarily and Functionally Unique, Specialised and Endangered), a composite conservation metric that captures both species’ evolutionary and functional diversity. EFUSE eliminates the need to assume the phylogenetic gambit by incorporating both forms of distinctiveness directly. Other metrics, such as ecoEDGE (22), have similarly integrated evolutionary and functional information, but EFUSE extends this approach by explicitly using a functional dissimilarity matrix and applying flexible weighting to the evolutionary, functional, and endangerment components. This ensures trade-offs between components are explicit and easily interpretable. By introducing EFUSE, we aim to advance our understanding of the links between ecological function, species resilience, evolutionary history, and conservation value, while better accounting for the diverse priorities and trade-offs inherent in conservation decision-making.

## Results

There was limited congruence between EDGE and FUSE scores for both mammals and birds (Fig. 1), highlighting that these metrics capture distinct aspects of biodiversity. To address this, we developed EFUSE, which integrates their complementary components (Fig. 2 and 3).

**Figure 1.**
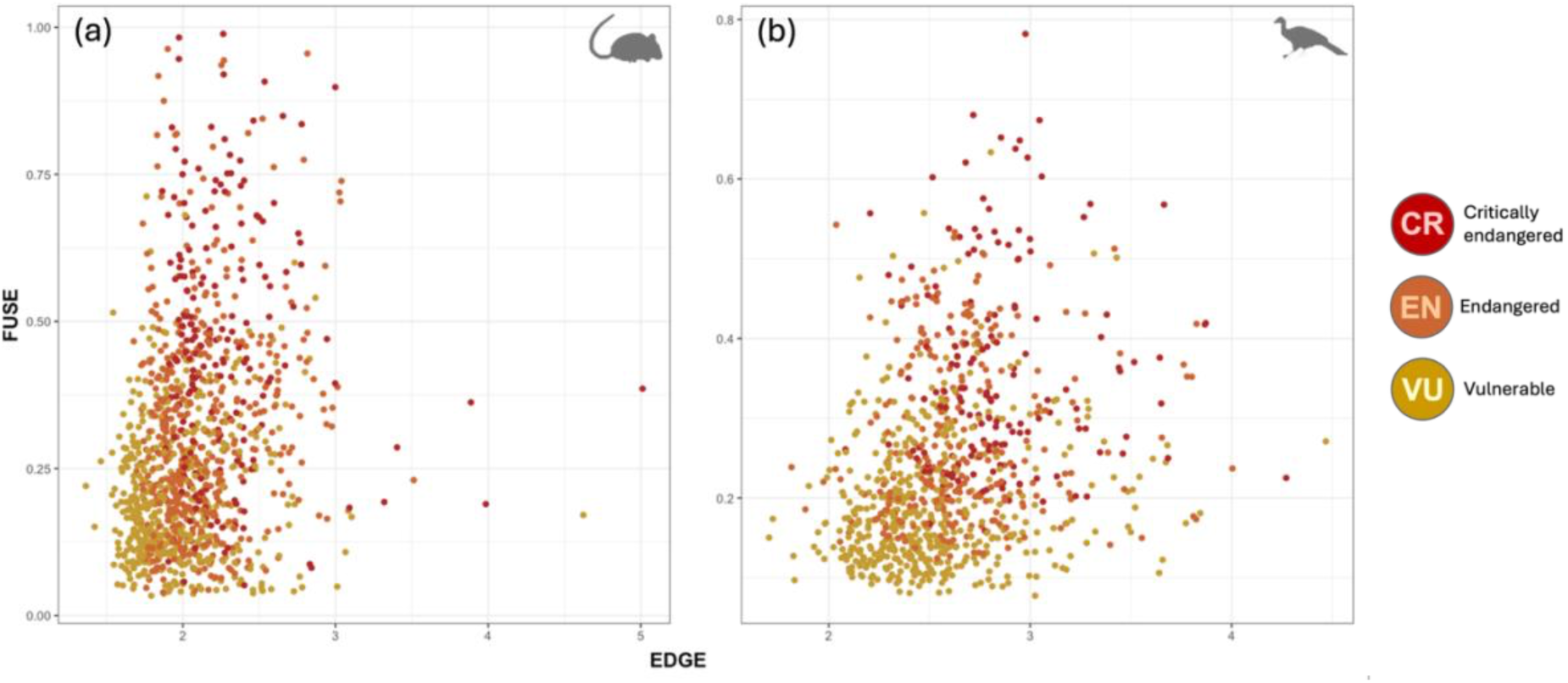
Plot of EDGE scores against FUSE score for (a) mammal and (b) bird species. Colours denote the global endangerment (GE) status of species according to the IUCN Red List of Threatened Species. Only species categorised as Vulnerable (VU), Endangered (EN) and Critically Endangered (CR) on the IUCN Red List are classed as EDGE species, near threatened (NT) and Least Concern (LC) species have therefore been excluded here. A linear regression was fitted to the data (EDGE ∼ FUSE) for mammals (F_1,4560_ =1103, r^2^ = 0.195, p < 0.0001) and birds (F_1,7893_ =1506, r^2^ = 0.160, p < 0.0001).

**Figure 2.**
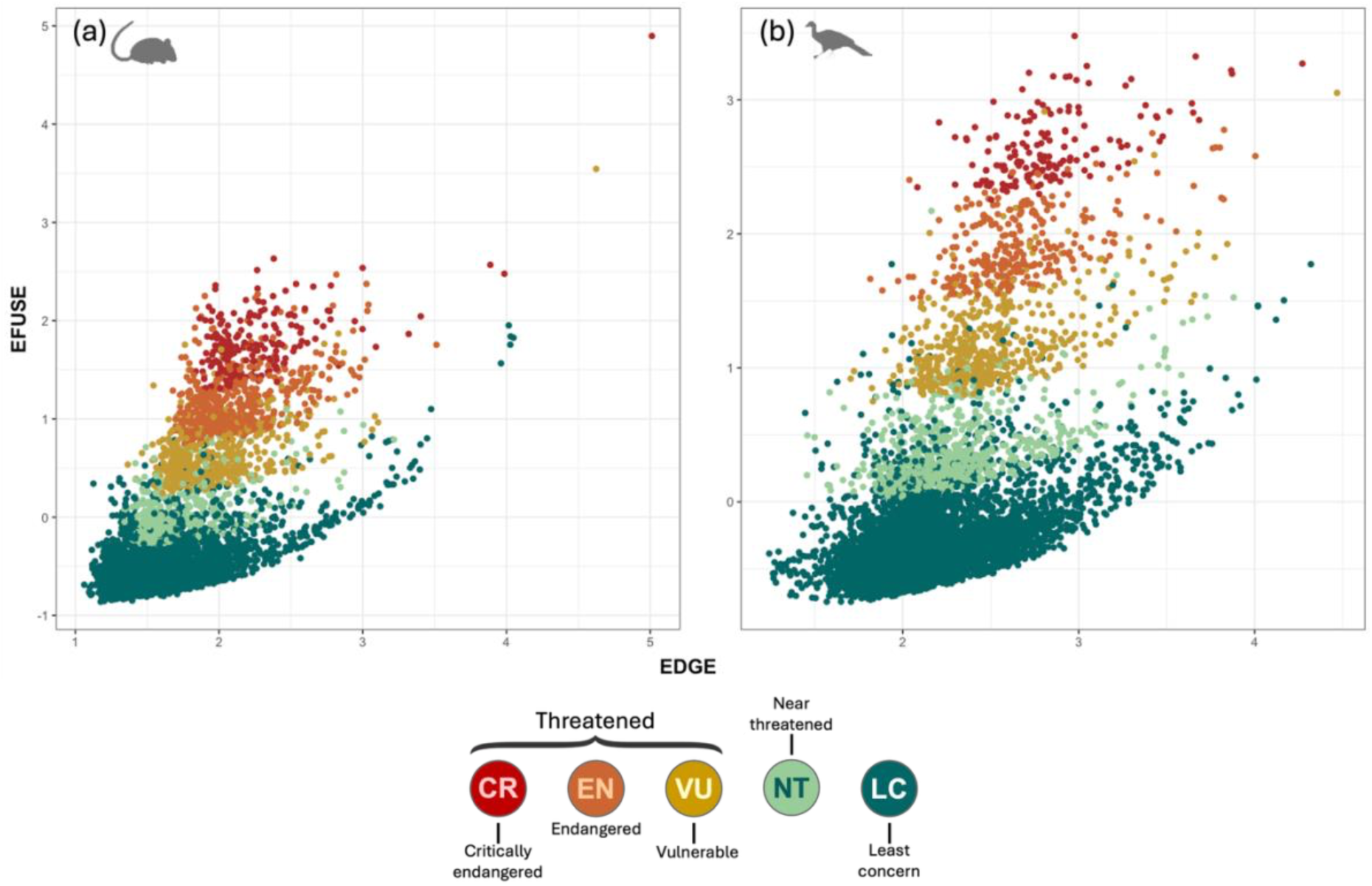
Plot of EDGE scores against EFUSE scores for (a) mammals and (b) birds. EFUSE incorporates the evolutionary distinctiveness (ED) and global endangerment (GE) status of the EDGE metric, in addition to the functional components of another conservation metric, FUSE. Colours denote the GE status of species according to the IUCN Red List of Threatened Species, where, from most to least endangered: Critically Endangered = CR; Endangered = EN; Vulnerable = VU; Near Threatened = NT; and LC = Least Concern. Note, species classed as NT and LC on the IUCN Red List are excluded from EDGE lists; they have been included here for comparison purposes.

**Figure 3.**
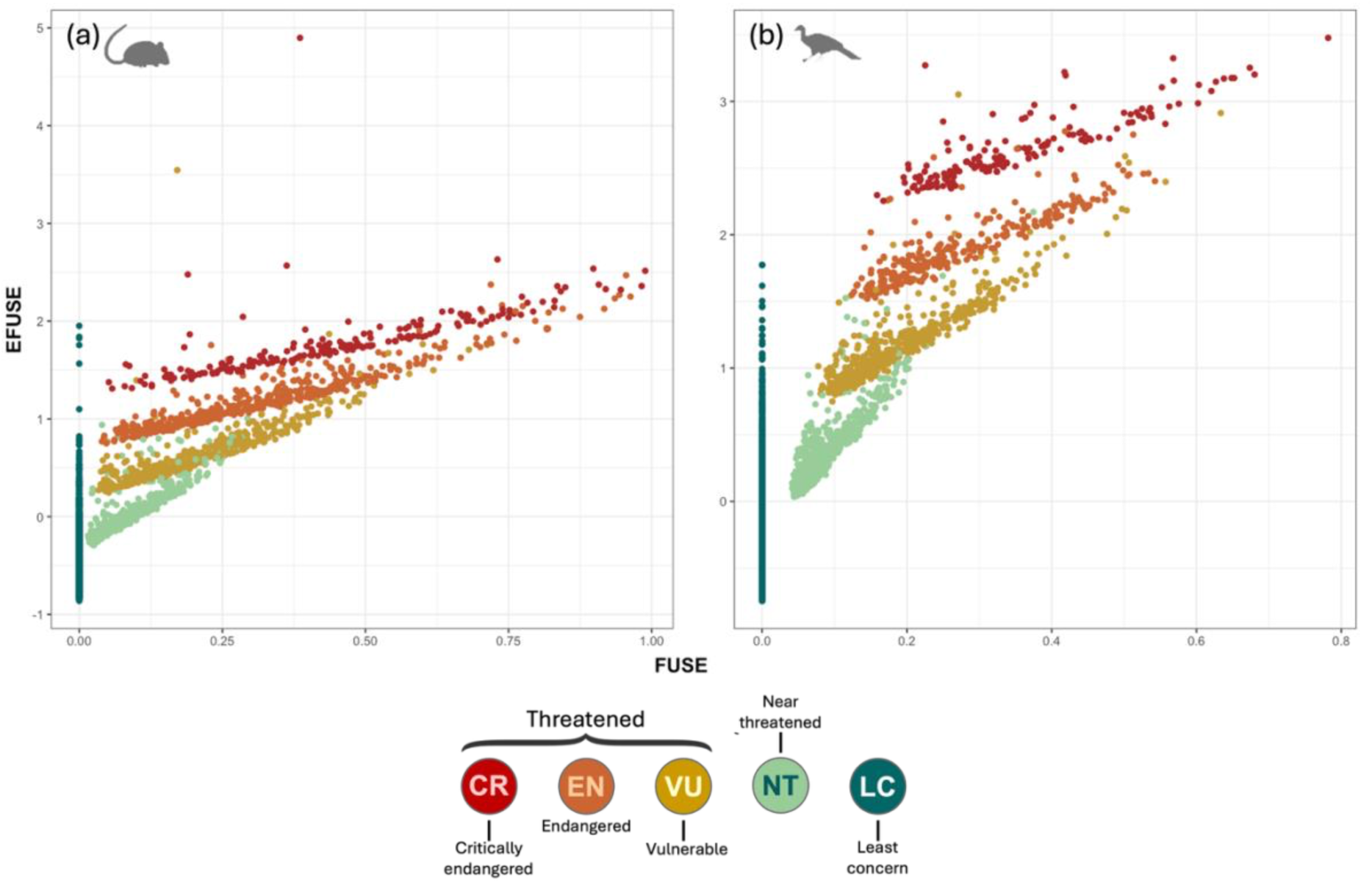
Plot of FUSE scores against EFUSE scores for (a) mammals and (b) birds. EFUSE incorporates the functional uniqueness (FUn), functional specialisation (FSp) and global endangerment (GE) status of the FUSE metric, in addition to the evolutionary component of another conservation metric, EDGE. Note different scale on both axes for (a) and (b). Colours denote the global endangerment (GE) status of species according to the IUCN Red List of Threatened Species, where, from most to least endangered: Critically Endangered = CR; Endangered = EN; Vulnerable = VU; Near Threatened = NT; and LC = Least Concern. Note, species classed as LC on the IUCN Red List always have a FUSE value of zero.

### Mammals

To establish how congruent EDGE and FUSE are, we compared the number of species within both the top 100 EDGE and top 100 FUSE species (Table S3). Only seven species are in both (Table S4). We found no significant correlation between ED, from EDGE, and the combined functional distinctiveness metric (𝐹𝑈𝑛 + 𝐹𝑆𝑝), from FUSE (Pearson’s *r* = 0.027, 95% confidence interval = –0.002, 0.056, *p* = 0.069).

EFUSE scores ranged from a minimum of -0.861 (Drab Atlantic tree-rat, *Phyllomys dasythrix*, LC on the IUCN Red List) to a maximum of 4.90 (Bolivian chinchilla rat, *Abrocoma boliviensis,* CR on the IUCN Red List; Table 2 & S2). The median EFUSE score for mammals was -0.359.

**Table 1.**
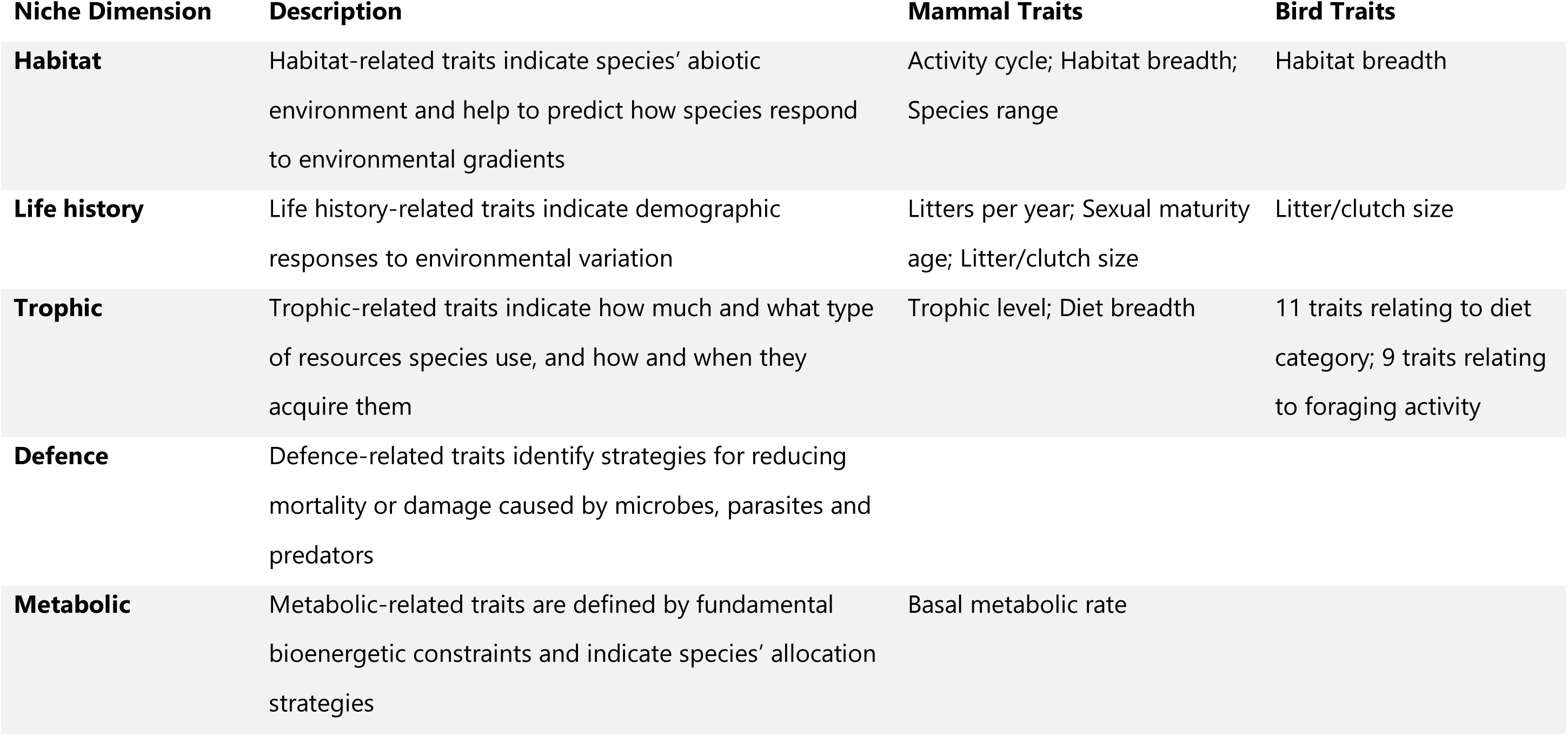
Descriptions of the five niche dimensions, each of which constitutes a fundamental aspect of a species’ niche (80, 83), and corresponding traits selected for use in the FUSE and EFUSE calculations. Body mass relates to several niche dimensions and was, therefore, also selected. Full trait descriptions and data sources are provided in Table S1 in the Supplementary Materials.

**Table 2.** List of 10 mammals with the highest EFUSE scores (the EFUSE mammal top-10). EDGE and FUSE rankings are from highest score (1) to lowest. For the EFUSE mammal top-100 see Table S2 in the Supplementary Materials. Note: EDGE and FUSE scores were recalculated in this study using the original EDGE1 methodology (7) for comparative purposes; therefore, species rankings may differ from those presented in the official EDGE lists.

| Species | Common name | Order | EFUSE Score | EDGE Rank | FUSE Rank |
| --- | --- | --- | --- | --- | --- |
| ***Abrocoma boliviensis*** | Bolivian chinchilla rat | Rodentia | 4.90 | 1 | 300 |
| ***Abrocoma bennettii*** | Bennett's chinchilla rat | Rodentia | 3.54 | 2 | 878 |
| ***Bradypus pygmaeus*** | Pygmy three-toed sloth | Pilosa | 2.63 | 331 | 41 |
| ***Tokudaia muenninki*** | Muennink's spiny rat | Rodentia | 2.57 | 9 | 342 |
| ***Hypogeomys antimena*** | Malagasy giant rat | Rodentia | 2.54 | 55 | 11 |
| ***Desmana moschata*** | Russian desman | Eulipotyphla | 2.51 | 455 | 1 |
| ***Zyzomys pedunculatus*** | Central rock rat | Rodentia | 2.48 | 7 | 789 |
| ***Neomonachus schauinslandi*** | Hawaiian monk seal | Carnivora | 2.47 | 89 | 4 |
| ***Macaca nigra*** | Celebes crested macaque | Primates | 2.37 | 216 | 10 |
| ***Addax nasomaculatus*** | Addax | Artiodactyla | 2.37 | 49 | 46 |

To establish the similarity between EDGE and EFUSE and FUSE and EFUSE we also compared these top-100 lists. 19 species were in both the EDGE and EFUSE top-100 lists and 76 species were in both the FUSE and EFUSE top-100 lists.

### Birds

Only six species are in both the EDGE top-100 birds and FUSE top-100 birds (Table S6 & S7). We found a weakly positive correlation between ED, from EDGE, and the combined functional distinctiveness metric (𝐹𝑈𝑛 + 𝐹𝑆𝑝), from FUSE (Pearson’s *r* = 0.060, 95% confidence interval = 0.038, 0.082, *p* < 0.001).

EFUSE scores ranged from a minimum of -0.745 (Spotted barbtail, *Premnoplex brunnescens*, LC on the IUCN Red List), to a maximum of 3.48 (Maleo, *Macrocephalon maleo*, CR on the IUCN Red List; Table 3 & S5). The median EFUSE score for birds was -0.338.

**Table 3.** List of 10 birds with the highest EFUSE scores (the EFUSE bird top-10). EDGE and FUSE rankings are from highest score (1) to lowest. For the EFUSE bird top-100 see Table S2 in the Supplementary Materials. Note: EDGE and FUSE scores were recalculated in this study using the original EDGE methodology (7) for comparative purposes; therefore, species rankings may differ from those presented in the official EDGE lists.

| Species | Common name | Order | EFUSE Score | EDGE Rank | FUSE Rank |
| --- | --- | --- | --- | --- | --- |
| ***Macrocephalon maleo*** | Maleo | Galliformes | 3.48 | 337 | 1 |
| ***Spheniscus demersus*** | African penguin | Sphenisciformes | 3.32 | 34 | 14 |
| ***Rowettia goughensis*** | Gough finch | Passeriformes | 3.27 | 3 | 445 |
| ***Otus insularis*** | Seychelles scops owl | Strigiformes | 3.25 | 280 | 3 |
| ***Anthracoceros montani*** | Sulu hornbill | Bucerotiformes | 3.22 | 15 | 91 |
| ***Mergus octosetaceus*** | Brazilian merganser | Anseriformes | 3.20 | 799 | 2 |
| ***Buteo ridgwayi*** | Ridgway's hawk | Accipitriformes | 3.19 | 14 | 89 |
| ***Gyps indicus*** | Indian vulture | Accipitriformes | 3.18 | 359 | 5 |
| ***Ardeotis nigriceps*** | Great Indian bustard | Otidiformes | 3.18 | 490 | 4 |
| ***Haliaeetus vociferoides*** | Madagascar fish eagle | Accipitriformes | 3.17 | 386 | 6 |

Comparison of the EDGE, FUSE and EFUSE top-100 lists revealed 20 species are on both the EDGE and EFUSE top-100 bird lists and 51 species are on both the FUSE and EFUSE top-100 bird lists.

### Phylogenetic dispersion

High EDGE scores were more dispersed in both the mammalian and avian phylogenies than high FUSE sores (Fig. 4).

**Figure 4.**
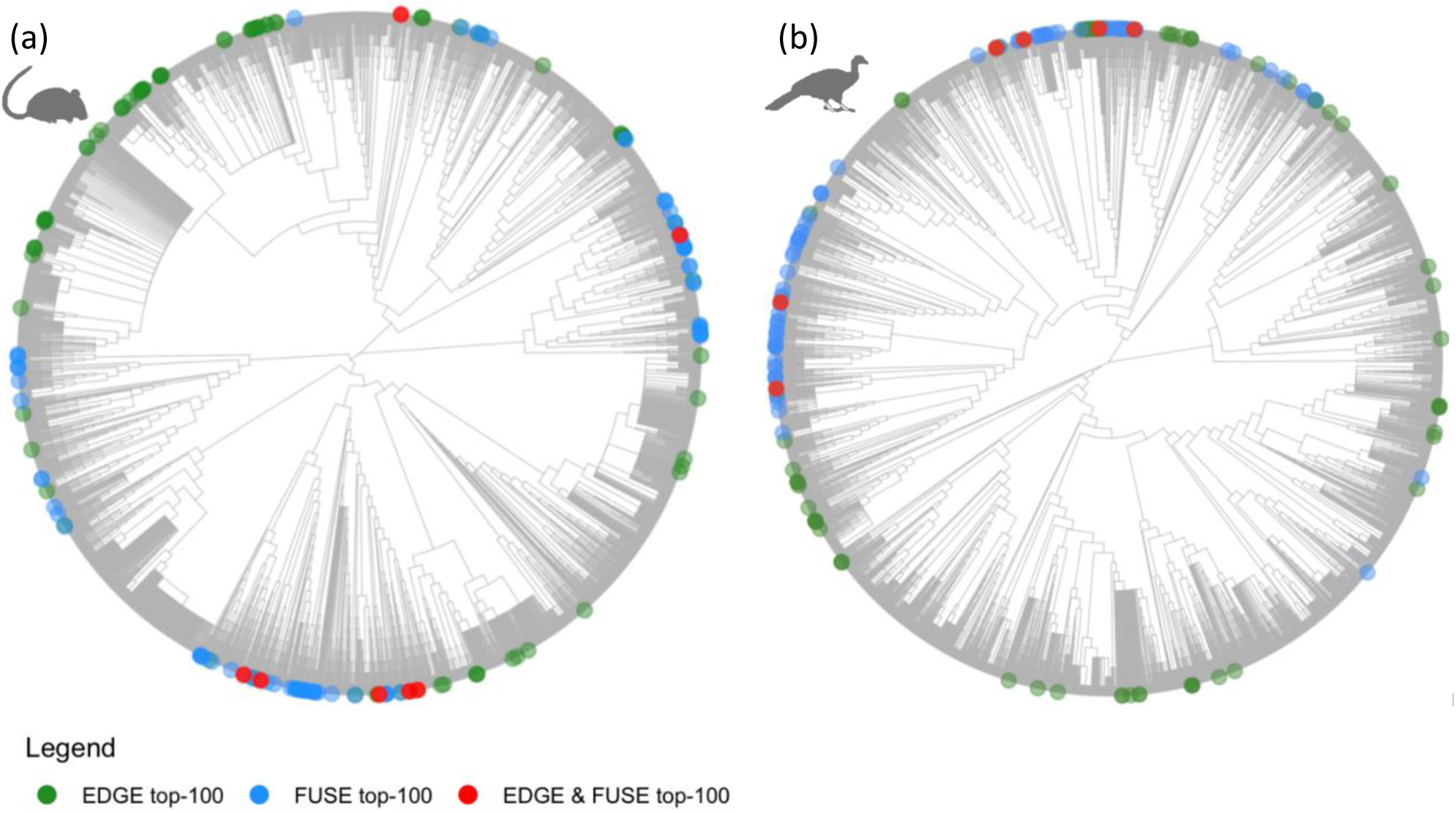
(a) Mammalian and (b) avian consensus phylogenies. Green circles indicate species in their respective top-100 EDGE list. Blue circles indicate species in their respective top-100 FUSE list. Red circles indicate species appearing in both top-100 EDGE and FUSE lists. The phylogenies shown represent 50% majority-rule consensus trees derived from 100 phylogenies for both mammals and birds. Across the 100 trees, the median D statistic indicated that species in the FUSE top-100 were more phylogenetically clustered (mammals = 0.544, P(random) < 0.001, P(Brownian) < 0.001, birds = 0.663, P(random) < 0.001, P(Brownian) < 0.001) than species in the EDGE top-100 (mammals = 0.688, P(random) < 0.001, P(Brownian) < 0.001; birds = 0.765, P(random) < 0.001, P(Brownian) < 0.001).

### Spatial congruence between EDGE and FUSE

Mean residuals from the regression of EDGE scores against FUSE scores varied substantially across space for both mammals and birds, indicating spatial heterogeneity in the congruence between the two metrics (Fig. 5).

**Figure 5.**
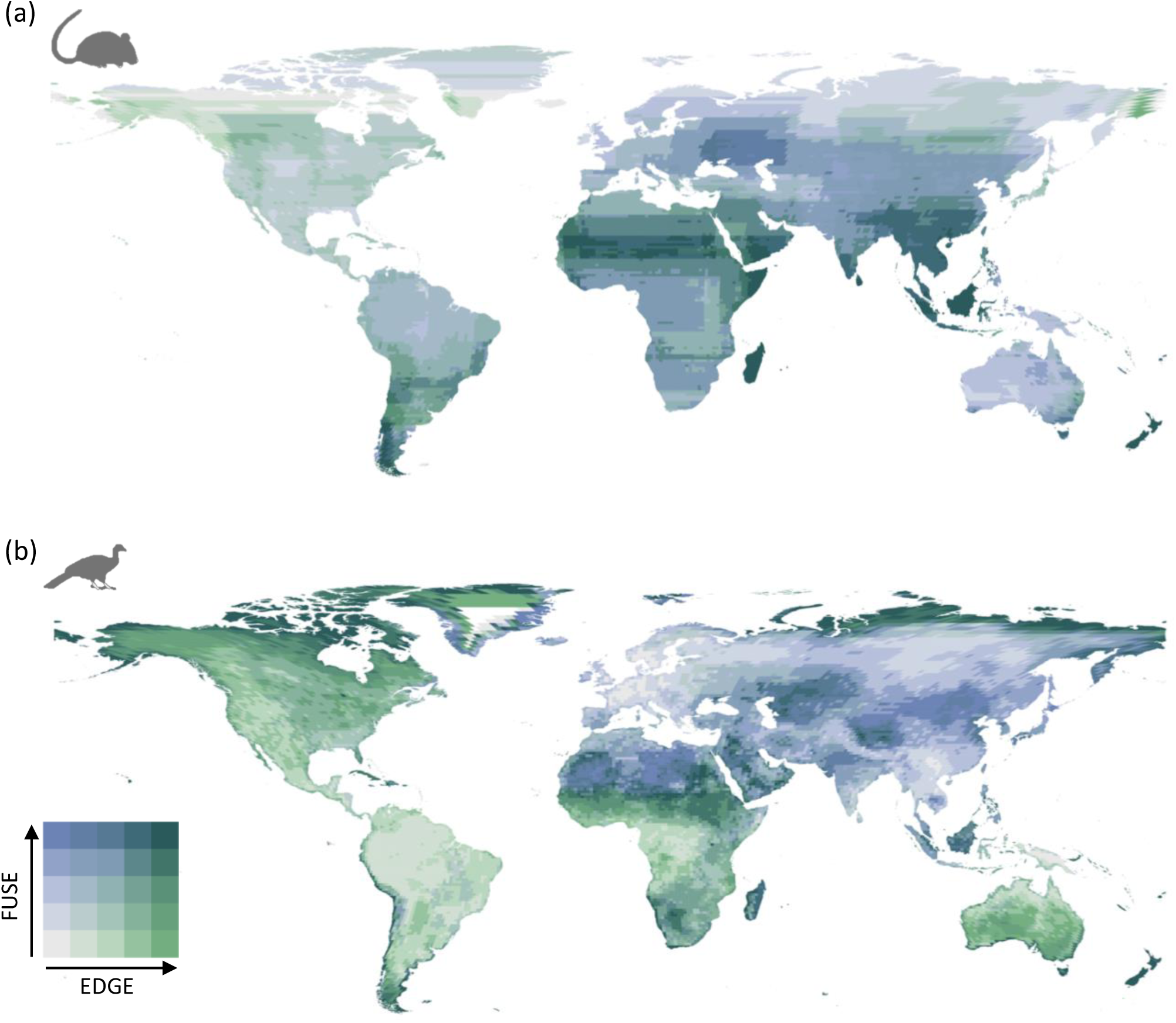
Global bivariate maps of the spatial distribution of EDGE and FUSE scores for (a) mammals and (b) birds. Green tones indicate areas of high EDGE scores, while blue tones indicate high FUSE scores. Teal tones indicate areas of both high EDGE and FUSE scores. Values were averaged within ∼100 km equal-area grid cells in a Mollweide projection. Figure generated using QGIS 3.42.1.

## Discussion

The lack of congruence between EDGE and FUSE reveals that the preferential application of one may result in species of high conservation value being inadequately protected. The EFUSE metric presented here provides a method for incorporating both evolutionary and functional components of biodiversity into conservation planning, thereby avoiding the potentially significant effects that disregarding either component may have on ecosystem functioning and stability and the maintenance of biodiversity.

### FUSE

To our knowledge, this is the first time that FUSE scores have been calculated for all extant mammals and birds, quantifying the conservation value of species according to their functional distinctiveness and endangerment status. The highest-ranking mammal, the Russian desman (*Desmana moschata*), and bird, the maleo (*Macrocephalon maleo*), exemplify how extreme ecological specialisation can elevate species under FUSE but not necessarily under EDGE. The Russian desman – a critically endangered, semiaquatic mole-like mammal confined to river basins in western Russia – has evolved specialised morphological and physiological traits for an aquatic lifestyle that are exceptionally rare among mammals and virtually unique within its clade (Talpidae) – a group that already exhibits some of the most specialised suites of morphological characters among small mammals (29, but see 30 for alternative phylogeny with Desmanines as a separate clade from Talpidae). Similarly, the maleo, a critically endangered megapode endemic to the Indonesian islands Sulawesi and Buton (31, 32), exhibits a highly distinctive reproductive strategy, burying its eggs in geothermal or sun-warmed sands and providing no parental care (33, 34). In both cases, unique ecological strategies rather than deep phylogenetic isolation drive their high FUSE scores relative to their EDGE rankings.

### Are EDGE and FUSE congruent?

For both mammals and birds, there was limited congruence between EDGE and FUSE scores (Fig. 1), as well as between ED, from EDGE, and the combined functional distinctiveness metric (𝐹𝑈𝑛 + 𝐹𝑆𝑝), from FUSE (Fig. S1). Notably, only seven mammal species (Table S4) and six bird species (Table S7) overlapped between the respective top-100 EDGE and FUSE lists. In addition, the top-100 mammals and birds with the highest FUSE scores were strongly clustered within their respective phylogenies (Fig. 4). Of the FUSE top-100 mammals, 67% belong to just three orders (Artiodactyla, Carnivora and Primates), despite these orders comprising just 17.1% of all mammal species included. Whilst phylogenetic overdispersion would be expected for species with high EDGE scores, given EDGE is designed to maximise unique branch length (7), the phylogenetic clustering of species with high FUSE scores is highly informative. It indicates that, consistent with the conditions Mazel, Mooers (21) proposed would reduce congruence between ED and FD, species within these clades display greater divergent evolution, with weaker or no stabilising selection, by comparison with other mammalian clades, leading them to be highly separated in functional space despite being closely related.

We also observed clear regional differences in the joint distribution of EDGE and FUSE scores. For mammals, congruence between the two metrics was strongest across tropical regions and several island systems, including New Zealand, where both evolutionary and functional distinctiveness are high (Fig. 5a), indicating that these areas contain assemblages where ancient lineages also occupy unique ecological roles. Areas of low congruence likely reflect cases where ED serves as a poor surrogate for FD, for which two main theories have been proposed (20). Firstly, in regions where ecologically distinct close relatives coexist, EDGE tends to prioritise only one of them, whereas both will likely be prioritised under FUSE. This pattern is evident in Patagonia, where the culpeo (*Lycalopex culpaeus*) cooccurs with the South American grey fox (*Lycalopex griseus*), a closely related but ecologically distinct relative. Secondly, in regions with strong niche conservatism, lineages will accumulate ED without corresponding increases in FD. Strong ecological niche conservatism is common in mammals (21, 35, 36) and will lead to highly evolutionarily distinct species being selected under EDGE, but not under FUSE as they will not be particularly functionally distinctive. This pattern typifies Australia, where mammals are evolutionarily isolated (37) but often retain conserved ecological roles (38), resulting in EDGE dominance.

For birds, congruence between EDGE and FUSE was highest in sub-Saharan Africa, New Zealand, and the far north of Canada and Russia, where both EDGE and FUSE scores are elevated (Fig. 5b). In contrast, low congruence occurred across much of Asia and northern Africa, where FUSE dominated, and parts of North America and Australia, where EDGE dominated. These regional differences likely reflect contrasting evolutionary dynamics across latitudes. In tropical regions, “basal” bird clades are disproportionately diverse and exhibit strong ecologically differentiation (Hawkins et al., 2007), leading ED and FD to covary closely, producing high congruence between EDGE and FUSE. This likely explains the high level of congruence observed for sub-Saharan Africa. In extratropical regions, rapid radiations and trait convergence within a few adaptive clades (e.g. passerines and waterfowl; 39, 40) have produced species that are functionally similar but differ little in ED – leading FUSE to prioritise ecological outliers, while EDGE highlights older, isolated lineages. Together, these spatial patterns demonstrate that the relationship between phylogenetic and functional diversity is not universal but contingent on lineage-specific histories of diversification and trait evolution, which vary strongly across latitude and clade.

The lack of congruence between the EDGE and FUSE lists reinforces concerns that ED is an imperfect surrogate for FD (20, 41, 42), with potentially substantial implications for conservation. The EDGE framework has been instrumental in elevating the role of phylogenetic diversity within conservation and underpins the EDGE of Existence programme (the programme now uses EDGE2 which we do not assess here), which guides numerous conservation projects and informs global biodiversity policy (14, 16, 43). By comparison, FUSE has not yet been applied in practical conservation contexts. As such, many species which are highly functionally unique and/or specialised are not being considered conservation priorities because it is assumed ED (and therefore their EDGE scores) adequately captures their FD (18), which our results show is not the case (Fig. S1). The phylogenetic clustering in FUSE further exemplifies this issue – many highly functionally distinct species in these clusters would not be prioritised by EDGE because they are closely related.

The potential loss of FD which could result from relying on EDGE – and therefore failing to adequately capture FD – has significant implications for ecosystem functioning. FD is now widely recognised as a critical biodiversity component which should be considered when devising conservation strategies (11, 24, 44, 45). High diversity of traits within a community allows species to fill diverse niches, exploit a diversity of resources, and efficiently assimilate energy and nutrients and transfer it within and across ecosystems. As such, high FD allows species to enhance and stabilise ecosystem processes (46–48). The loss of FD which could result from failing to adequately quantify, consider and, thus, protect it could destabilise ecosystem processes and affect ecosystem functioning over large scales (49, 50).

### Why does evolutionary distinctiveness matter?

Given the limited congruence between EDGE and FUSE rankings for mammals and birds, we must consider the implications of choosing one over the other as a guide for conservation planning. Importantly, we should not disregard the benefits conserving high-priority EDGE species may yield. Proponents of EDGE do not merely encourage the prioritisation of ED as a potential surrogate for FD, they believe maximising ED serves more broadly to conserve a wide variety in forms and functions and thus potential future utilitarian value (18, 45). The underlying rationale is that phylogeny captures both known and unknown features of organisms (45, 51, 52). In maximising ED, and, therefore, maximising diversity in these features, EDGE proponents believe it also serves to maximise option value, which is the value of biodiversity derived through the provision of benefits and uses, often unanticipated, for future generations (18). In this view, greater biodiversity is seen to represent greater option value (53). However, the lack of congruence between EDGE and FUSE, as shown here, and ED and FD, as demonstrated previously (20, 21, 54), calls into question the ability of ED to capture diversity in forms and functions, and, therefore, to truly optimise option value.

Further, the nature of option value makes it difficult to quantify and thus to assess the ability of PD to capture it. Yet, Faith (55) would argue that option value can never be directly quantified and, therefore, assessed, because option value is, by definition, not something we know about yet.

Beyond option value, a species’ unique contribution to ED is also seen as a proxy for ’evolutionary novelties’, which relate to bequest and existence values (45). Whilst the importance of such intrinsic values of species is controversial academically (e.g. 56, 57-59), it is undeniable that these values are extremely important to many cultures and individuals (60, 61). As such, the ability of ED to capture bequest and existence values is highly valuable.

Additionally, ED safeguards lineages that embody vast spans of evolutionary history. Such species represent irreplaceable records of the diversification of life, even if their present-day ecological roles are modest.

### The best of both - EFUSE

EDGE and FUSE both incorporate species attributes that the other does not. EDGE captures evolutionary history, option value, bequest, and existence values; FUSE captures functional diversity. These two approaches should be complementary – they are based on different components of biodiversity and provide distinct information on the conservation value of species. Here, we present a new metric, EFUSE, that consolidates the key components of both EDGE and FUSE to streamline their incorporation into conservation decision-making. The purpose of EFUSE is to simplify the number of components which must be considered when planning conservation, consolidating the ED, FD and GE of species into one score which can be directly compared between species or averaged across communities, geographical areas and phylogenetic clades.

EFUSE is not the first metric to include both evolutionary and functional components; Hidasi-Neto, Loyola (22) proposed ‘ecoEDGE’ as an approach to build upon EDGE by incorporating species’ functional distinctiveness. EFUSE differs from ecoEDGE in two distinct ways. Firstly, the functional component of ecoEDGE (‘EcoD’) is calculated using a functional dendrogram, whereas EFUSE uses a trait-based dissimilarity matrix, as per Pimiento, Leprieur (9). This reflects a conceptual distinction: while phylogenetic depictions of evolution are not always appropriate (62), this is arguably less controversial than forcing all functional traits into a hierarchical structure. In some cases, dendrogram-based approaches can distort trait relationships (63, 64). By instead using a dissimilarity matrix, EFUSE avoids this issue and more directly reflects the structure of the underlying trait data (65, 66). Secondly, unlike in EFUSE, a weighting factor is not applied to GE (see supplementary materials for full equation), precluding the option to increase or lessen endangerment’s contribution to species’ scores. In addition, the use of single weighting factors for the functional and evolutionary components (𝛼) and combined functional and evolutionary component and GE (𝛽) in EFUSE means the compromises made between phylogenetic and functional information and GE are made explicit and easy to interpret.

The mammal species with the highest EFUSE score was the Bolivian chinchilla rat (*Abrocoma boliviensis*), ranked as CR on the IUCN Red List. This species is endemic to a highly restricted region of the Andes in Bolivia (67, 68) and belongs to the family Abrocomidae, which are estimated to have diverged at ∼30–40 Ma (69). This deep evolutionary split, combined with its limited geographic range, likely underpins the species’ high EFUSE score. Despite its apparent conservation importance, remarkably little is known about its current population status or ecological role (70). The bird species with the highest EFUSE score was the Maleo (*Macrocephalon maleo*), ranked as CR on the IUCN Red List. Like A. boliviensis, it remains poorly studied, although it is known to be threatened by a suite of anthropogenic pressures, including egg poaching and habitat fragmentation (71). Notably, the Maleo was also identified as the bird species with the highest FUSE score. As discussed previously, the Maleo exhibits a highly distinctive reproductive strategy, making it functionally distinctive among birds. In addition, the Maleo is the sole member of the *Macrocephalon* genus, contributing to its high evolutionary distinctiveness with few close relatives. Together, these examples illustrate EFUSE’s potential to highlight species that are both evolutionarily and functionally distinct, but which remain relatively understudied and disproportionately overlooked in conservation.

As already mentioned, we used original EDGE methodology (7) to calculate EDGE scores for comparison with FUSE, and to calculate the ED component of EFUSE. This was appropriate for examining discrepancies between ED-based and FD-based prioritisation, and for guiding the development of EFUSE, which does not merely assume, but directly tests and is informed by the phylogenetic gambit. Future work could explore incorporating PD complementarity into the ED component, following advances made in EDGE2 (17) and HEDGE (a mathematically equivalent metric to EDGE2; 72). By accounting for the extinction risk of close relatives, PD complementarity better reflects a species’ expected future contribution to PD. Incorporating such approaches alongside functional information could provide a more accurate and forward-looking assessment of species’ irreplaceability (73, 74). Here, we have integrated functional trait information into the original EDGE framework through EFUSE. A natural next step would be to extend this approach to EDGE2, incorporating functional traits, for example via a functional dissimilarity matrix or traitgram, alongside PD complementarity to create an even more comprehensive, phylogenetically and functionally informed prioritisation metric. We emphasise, however, that just because something *can* be done, does not mean that it *must* be done, and that a conservation program should only make use of a metric if it wishes to, and if the metric is compatible with its goals and underlying narrative.

### Conclusion

The limited congruence between EDGE and FUSE is alarming. EDGE is the most commonly applied metric in conservation planning, and yet, via direct empirical comparison, we demonstrate it largely fails to capture species with high functional distinctiveness (i.e., high FUSE species). Given components of both EDGE and FUSE are complementary and capture information on important species attributes which relate to ecosystem functioning, it has become necessary to combine their components into a single, versatile metric. The metric we present here – EFUSE – simplifies the number of components to consider in conservation planning and is highly adaptable to specific stakeholder values and purposes.

Our findings demonstrate the importance of examining the potential implications of subjective conservation decisions. EFUSE removes the need to choose between valuing evolutionary and functional distinctiveness and thereby helps to facilitate the incorporation of information on more important biodiversity attributes into conservation decision-making. Future uses of EFUSE should include the use of extinction scenarios to estimate extinction probability given a certain duration (e.g., 100 years) of status quo conservation (e.g. following 75). This would allow us to forecast declines in evolutionarily and functionally distinct species in order to better understand the scale and impact of species loss. In combination with the EFUSE lists presented here, this will help to identify highly vulnerable and important species for which we should particularly focus conservation efforts in order to preserve ecosystem functions and stability, and option, bequest and existence values.

## Methods

To assess the congruence between the EDGE and FUSE matrices, we calculated EDGE and FUSE scores independently for mammals and birds. These groups were selected because comprehensive evolutionary and functional trait data are readily available for a large proportion of species (76–78) . We also developed and calculated a new metric – EFUSE (Evolutionarily and Functionally Unique, Specialised, and Endangered) – which integrates components of both EDGE and FUSE, aiming to provide a more comprehensive prioritisation framework for conservation. The EDGE, FUSE, and EFUSE indices are all relative measures that combine species’ distinctiveness with their level of endangerment. A high score in any of the three indicates that a species represents a disproportionate amount of evolutionary or functional distinctiveness – or both – and is also highly threatened. All analyses were conducted using R version 4.4.1 (79).

### Data and Trait Estimation

We compiled phylogenetic and ecological trait for 5319 mammal species and 7932 bird species. We selected traits that correspond to the ‘five niche dimensions’ proposed by Winemiller et al. (80; Table 1). We selected these traits as they correlate to species’ environmental niches and, therefore, to their ecological role and contributions to ecosystem functioning (81, 82). Body size is integral to multiple dimensions (80) and was therefore also included.

Trait data were obtained from three major databases: a global mammal and bird functional trait database (78), a global species-level dataset of mammalian life-history, ecological and geographical traits (76), and an amniote life-history database (77). Habitat breadth was coded using the International Union for Conservation of Nature (IUCN) Habitats Classification Scheme and was quantified as the number of suitable habitats listed for each species (82). Due to limited data availability, it was not possible to include an even spread of traits from each niche dimension. To address this, we applied a weighting to each trait based on how many other traits corresponded to the same niche dimension. This ensured that traits from each dimension contributed equally (one part) to the species dissimilarity matrices (see ‘*Calculating FUSE scores’* for details on creating the matrices). Body size was treated as a separate component, contributing one equal part, given its broad relevance to multiple niche dimensions.

Trait data is still relatively limited for several traits and taxa (20). As such, we only selected traits for which 50% of species in a given clade had data available. For species with missing trait data, we imputed trait values using phylogenetic ancestral state estimation using the ‘phyestimate’ function in the *picante* package (84). To account for phylogenetic uncertainty, we repeated this process across 100 phylogenetic trees for each clade. These trees were randomly drawn from a pool of 1000 composite ‘supertree’ phylogenies for both mammals (species : nodes ratio >0.999; 85) and birds (species : nodes ratio >0.999; 86). We chose to select 100 phylogenies as a balance between computational feasibility and adequate representation of phylogenetic uncertainty; previous work suggests that relatively few trees (∼50) can be sufficient for robust inference (87). Imputation is imperfect and may introduce biases, given it is based on phylogenetic information (88). However, as we are testing the congruence of EDGE (phylogeny-based) and FUSE (trait-based), this bias should, in fact, make our findings of a lack of congruence conservative, the differences perhaps being even greater. EDGE, FUSE and EFUSE scores were also calculated across the same 100 phylogenetic trees for each clade. The median score for each metric across all trees was then taken as the final score for each species, reducing the influence of phylogenetic uncertainty on score estimation.

Following both EDGE and FUSE, we classified the GE value for all species based on their most recent classification on the IUCN (89). Species were assigned as follows: LC (Least Concern) = 0; NT (Near Threatened) = 1; VU (Vulnerable) =2; EN (Endangered) = 3; and CR (Critically Endangered) = 4. Data deficient species were not classified and therefore excluded from the study.

### Calculating EDGE scores

ED scores were calculated as described in Isaac, Turvey (7): sharing each phylogenetic branch equally among all its subtending species (Fig. 6(a)) such that ED is measured in units of time. ED scores were calculated using the ‘ed.calc’ function in the caper package (90).

**Figure 6.**
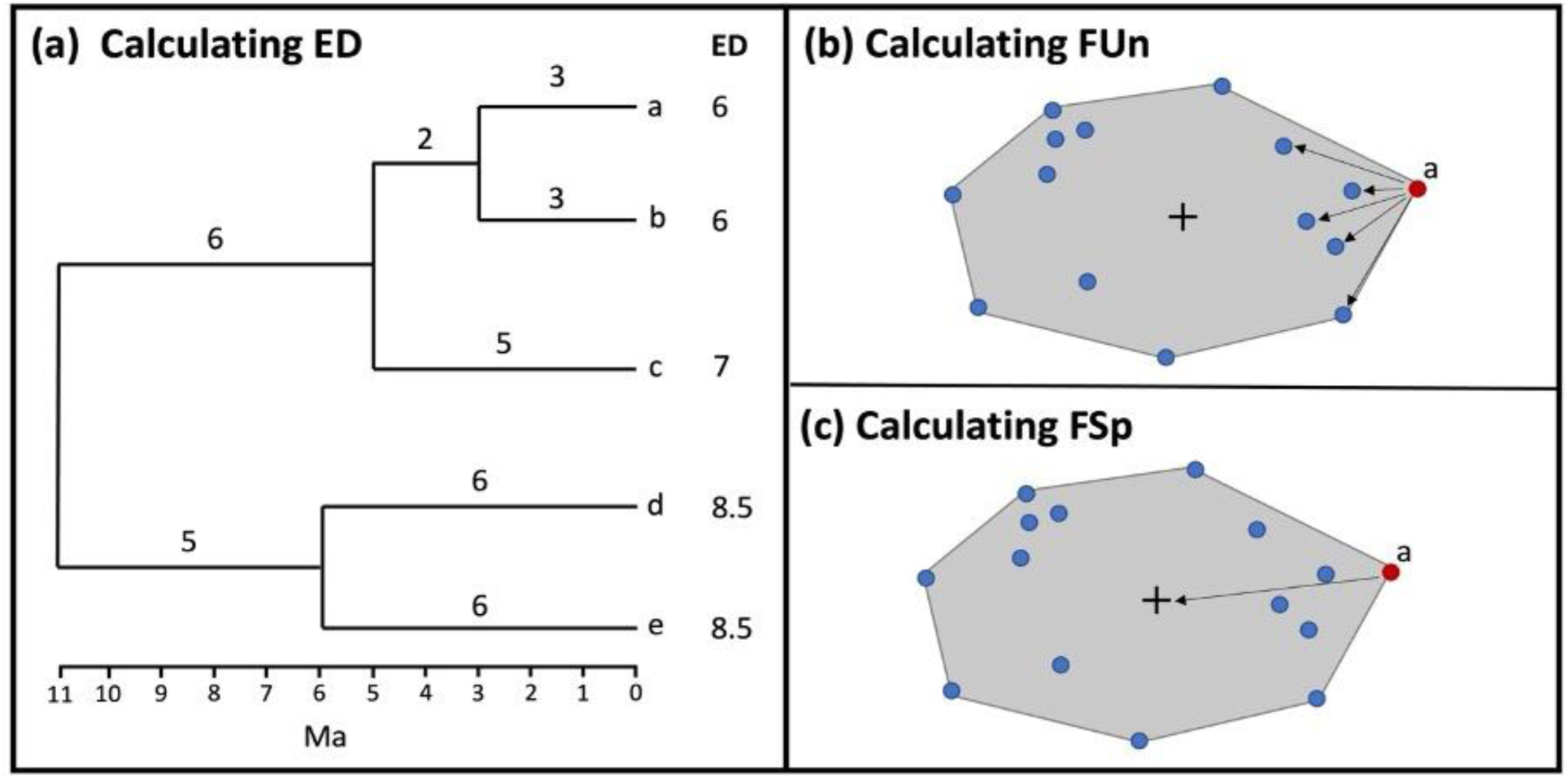
Schematics illustrating how to calculate evolutionary and functional measures. (a) displays a hypothetical phylogeny of five species with Evolutionary Distinctiveness (ED) scores. Numbers above each branch indicate the length in millions of years before present (Ma). The ED score of species a is given by the sum of the ED scores for each of the three branches between a and the root. The terminal branch contains just one species ‘a’ and is 3 million years (Ma) long, so receives a score of 3 Ma. The next branch is 2 Ma and contains two species, so each daughter species (a and b) receives 1 Ma. The next branch is 6 Ma and contains three species, so each daughter species (a, b and c) receives 2 Ma, so the total ED score for species a is given by (3/1+2/2+6/3)=6 Ma. (b) and (c) display a multidimensional ‘functional space’, in (b) the functional uniqueness (FUn) of species a is measured as its mean distance to a set of n nearest neighbours (here n = 5), characterising a’s local relative position in trait space. In (c) the functional specialisation (FSp) of species a is measured as its distance from the trait space centroid.

EDGE scores are calculated by summing the species’ ED scores and their IUCN Red List status (7) as follows:

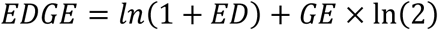

It is important to note that EDGE scores were calculated following the original EDGE (EDGE1) methodology (7). EDGE scores published by the EDGE of Existence Programme at ZSL have used the EDGE2 protocol since 2023 (17). The EDGE2 protocol includes several advances from the original EDGE metric including the incorporation of methods for dealing with uncertainty in the extinction risk of closely related species, and of PD complementarity into the ED component. Specifically, the extinction risk of a species’ close relatives is considered when calculating the ED component to better reflect the expected contribution of the species to PD in the future. Whilst this update has led to changes in published EDGE lists, it does not change the underlying assumption that the use of ED within the EDGE methodology adequately captures the functional distinctiveness of species. Thus, our use of the original, simpler EDGE1 formulation is appropriate for examining discrepancies between ED-based and FD-based prioritisation, and for guiding the development of future metrics that do not merely assume, but directly test and are informed by the phylogenetic gambit. In the case that phylogenetic and functional information provide identical information, our approach is robust, and in the case that they do not, our approach makes the impacts of the differences explicit.

### Calculating FUSE scores

FUSE scores were calculated as per Pimiento, Leprieur (9). For each clade, we created a species traits distance matrix using a modified version of Gower’s distance (’daisy’ function in the cluster package; 91). We used Gower’s distance because it can handle multiple data types and the inclusion of variable weights. From this functional dissimilarity matrix, we built a multidimensional Euclidean space based on a principal coordinates analysis (PCoA; 92) as a PCoA can be applied to distance matrices. We retrieved the PCoA axes using the ‘dudi.pco’ function of the *ade4* R package (93–97).

Using the multidimensional trait spaces, we calculated functional richness (herein, FRic), which measures the volume of functional space occupied, i.e., the convex hull volume whose vertices are delimited by the species at the edge of multidimensional trait space (98).

Functional uniqueness (FUn) sensu measures the level of isolation of each species inside the functional space (99), which allows for quantifying the level of species’ uniqueness or redundancy (100). For each clade, we calculated mean FUn considering the five nearest neighbours (Fig. 6(b)). Pimiento, Leprieur (9) found a strong correlation between FUn using five neighbours and using one, three and 10, demonstrating this metric for FUn to be robust to the number of neighbours used. We also calculated functional specialisation (FSp) as the Euclidean distance of each species to the centre of the multidimensional trait space (Fig. 1(c); 99), which allows distinguishing between those close to the centre of the space (displaying average trait combinations, or generalists) and those near the edges of the space (displaying extreme trait combinations, or specialists).

Several alternative formulations of FUSE are possible: FUSE’, which can be interpreted as the mean of FUn and FSp weighted by GE; and FUSE’’ which is the sum of FUn and FSp weighted by GE. The alternative formulations of FUSE have proven almost indistinguishable (101). As such, we follow the original formulation proposed in Pimiento, Leprieur (9), whereby FUSE is calculated by adding the product between species’ uniqueness and specialisation scores and their IUCN Red List status as follows:

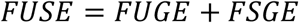

Where

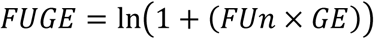

and

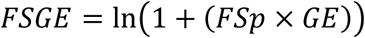

FUn and FSp are standardised between 0 and 1 and multiplied by four, which ensures they contribute equally to FUGE and FSGE as GE (which ranges between 0 and 4).

### Comparison of EDGE and FUSE

To assess the congruence between EDGE and FUSE, we performed a linear regression of EDGE scores on FUSE scores. Separate regressions were conducted for mammals and birds. Because extinction risk (GE) was classified using species’ IUCN Red List classifications for both EDGE and FUSE, introducing potential correlation, we also examined the relationship between evolutionary and functional distinctiveness independent of extinction risk (GE). We conducted a Pearson’s product-moment correlation test between ED (from EDGE) and a combined metric of functional uniqueness and specialisation (from FUSE), calculated as the sum of FUn and FSp.

To better understand how EDGE and FUSE scores relate to species’ positions in the phylogeny, we assessed whether species with high scores under each metric were phylogenetically clustered. If species with high EDGE scores are phylogenetically overdispersed while those with high FUSE scores are clustered, this would support the idea that the two metrics capture different aspects of biodiversity. Additionally, if species with high FUSE scores are phylogenetically clustered, this would indicate that functionally distinct species tend to occur within particular clades, highlighting patterns of functional divergence within phylogenetically related clades. To test this, we calculated phylogenetic *D* statistics (102) for the top 100 EDGE and FUSE mammal and bird species. We chose this quantity because the top-100 EDGE species are commonly referred to as the species which are the highest priority (7, 103). Unlike most diagnostics of phylogenetic signal, the *D* statistic can deal with categorical data, as required here. For each class, we assigned species two binary scores – one depending on whether they were one of the 100 species with the highest EDGE scores and one depending on whether they were one of the 100 species with the highest FUSE scores. Using the ‘phylo.d’ function in the *caper* R package, we then calculated the *D* value for each of these scores. The median D value and associated P-values across the 100 phylogenies were then taken as our summary estimates. A high *D* value indicates phylogenetic overdispersion, while low values indicate clustering. Because EDGE prioritises phylogenetic uniqueness by design, we expect those species to be more clustered. Clustering among high FUSE species; however, would reveal how functionally distinct species are distributed phylogenetically – and whether FUSE overlaps with or diverges from EDGE in the structure of species it prioritises.

ED has previously been shown to vary in how well it serves as a proxy for FD across geographical space (Mazel et al., 2018). It, therefore, seemed probable that the level of congruence between EDGE and FUSE would also vary spatially. To examine this hypothesis, we plotted the mean residuals from a linear regression of EDGE against FUSE. Residuals were calculated per species and then spatially aggregated across a global equal-area grid with a resolution of 1° latitude by 1° longitude (approximately 110 km × 110 km at the equator).

The resulting map visualises the spatial pattern of congruence between EDGE and FUSE, with low residual values indicating stronger agreement between the two metrics and higher values reflecting greater disparity.

### EFUSE

To create a more comprehensive metric, we built upon the complementary aspects of EDGE and FUSE. EFUSE incorporates the ED component of EDGE, the FUn and FSp components of FUSE, and the GE components of them both, as follows:

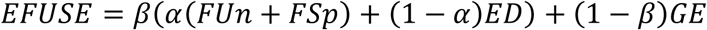

where ED is evolutionary distinctiveness (calculated as in the original EDGE methodology), FUn is functional uniqueness, and FSp is functional specialisation (calculated as in FUSE). 𝛽 and 𝛼 are weighting parameters which can range from 0 to 1. When 𝛽 = 1, EFUSE scores only incorporate evolutionary and functional components, not endangerment (GE). When 𝛽 = 0, EFUSE is only the level of endangerment. When 𝛼 = 1, EFUSE does not incorporate evolutionary distinctiveness; when 𝛼 = 0, EFUSE does not incorporate functional distinctiveness. When 𝛽 = 2⁄3 and 𝛼 = 1⁄2, as used here for mammal and bird EFUSE scores, the functional, evolutionary and endangerment components contribute equally.

This formulation makes EFUSE both conceptually powerful and practically flexible. The 𝛼 parameter explicitly governs the balance between evolutionary and functional distinctiveness, allowing us to test the core assumptions of the phylogenetic gambit. The 𝛽 parameter separates the prioritisation of species based on their intrinsic properties from their level of endangerment (GE), providing a way to examine how conservation priorities shift under different value systems or decision-making frameworks. For example, increasing α, and thereby making EFUSE more analogous to FUSE, would prioritise species with distinct ecological traits, such as functionally specialised pollinators or apex predators. Decreasing α, making EFUSE more analogous to EDGE, would shift focus towards evolutionarily unique species, thereby preserving the tree of life. Similarly, lowering 𝛽 would give greater weight to extinction risk and immediate conservation urgency, while higher 𝛽 values emphasise the long-term preservation of evolutionary and functional diversity. This structure not only enables hypothesis testing, but also ensures that EFUSE remains robust to uncertainty in trait data or threat classification. By making the trade-offs between the different components explicit, EFUSE offers a transparent and adaptable tool for conservation planning.

Each component in the EFUSE metric (𝐹𝑈𝑛, 𝐹𝑆𝑝, 𝐸𝐷 𝑎𝑛𝑑 𝐺𝐸) is scaled such that their means are zero and standard deviations are one. The sum of 𝐹𝑈𝑛 and 𝐹𝑆𝑝 was also scaled to ensure the functional component (𝐹𝑈𝑛 + 𝐹𝑆𝑝) and evolutionary component (𝐸𝐷) contribute equally to the measure before weighting parameters are applied, as was the sum of the scaled functional component (𝐹𝑈𝑛 + 𝐹𝑆𝑝) and scaled evolutionary component (𝐸𝐷). This ensures that each component contributes equally to the measure and that judgments about which components are more important are made readily explicit when utilising the weighting parameters (104).

A key distinction between EFUSE and existing metrics (EDGE and FUSE) is how species with low extinction risk are treated. Under EDGE, species listed as NT or LC are excluded, while in FUSE, species listed as LC always receive a FUSE score of zero. In contrast, EFUSE can assign non-zero values to such species if they are evolutionarily and/or functionally exceptional.

This occurs because EFUSE incorporates extinction risk as one component within a weighted additive framework, rather than applying it as a multiplicative constraint. The weighting parameter (𝛽) also provides flexibility to reduce or remove the endangerment component entirely. This allows EFUSE to highlight species that, although not currently threatened, make disproportionately large contributions to evolutionary history or ecosystem functioning. It has long been recognised that conservation tools must also be able to identify species of high evolutionary and ecological value before their decline begins, in order to maximise conservation efficiency and safeguard irreplaceable biodiversity before it reaches critical thresholds (105). EFUSE provides a framework that enables this flexibility.

### Data and code availability

All code supporting the analyses in this study are provided in support of this manuscript: https://github.com/lucysom/EFUSE.git.

## Supporting information

Supplementary Results & Discussion

## References

1. Ceballos G, Ehrlich PR, Barnosky AD, García A, Pringle RM, Palmer TM. Accelerated modern human–induced species losses: Entering the sixth mass extinction. Science Advances. 2015;1(5):e1400253.

2. Hull PM, Darroch SAF, Erwin DH. Rarity in mass extinctions and the future of ecosystems. Nature. 2015;528(7582):345–51.

3. Pimm SL, Jenkins CN, Abell R, Brooks TM, Gittleman JL, Joppa LN, et al. The biodiversity of species and their rates of extinction, distribution, and protection. Science. 2014;344(6187):1246752.

4. Bottrill MC, Joseph LN, Carwardine J, Bode M, Cook C, Game ET, et al. Is conservation triage just smart decision making? Trends in Ecology & Evolution. 2008;23(12):649–54.

5. Vane-Wright RI, Humphries CJ, Williams PH. What to protect?—Systematics and the agony of choice. Biological Conservation. 1991;55(3):235–54.

6. Baselga A, Martín-Devasa R, Gómez-Rodríguez C. Areas of High Biodiversity Value Evidenced by the Spatial Scaling of Phylogenetic Uniqueness. Ecology Letters. 2025;28(7):e70179.

7. Isaac NJB, Turvey ST, Collen B, Waterman C, Baillie JEM. Mammals on the EDGE: Conservation Priorities Based on Threat and Phylogeny. PLoS ONE. 2007;3:98.

8. Mace GM, Possingham HP, Leader-Williams N. Prioritizing choices in conservation. In: Macdonald D, Service K, editors. Key topics in conservation biology. Oxford, UK: Blackwell Publishing; 2006. p. 17-34.

9. Pimiento C, Leprieur F, Silvestro D, Lefcheck JS, Albouy C, Rasher DB, et al. Functional diversity of marine megafauna in the Anthropocene. Science Advances. 2020;6(16):eaay7650.

10. Isaac NJB, Pearse WD. The Use of EDGE (Evolutionary Distinct Globally Endangered) and EDGE-Like Metrics to Evaluate Taxa for Conservation. In: Scherson R, Faith DP, editors. Phylogenetic Diversity: Cham, Springer; 2018. p. 27-39.

11. Cadotte MW, Carscadden K, Mirotchnick N. Beyond species: functional diversity and the maintenance of ecological processes and services. Journal of Applied Ecology. 2011;48:1079–87.

12. Forest F, Grenyer R, Rouget M, Davies TJ, Cowling RM, Faith DP, et al. Preserving the evolutionary potential of floras in biodiversity hotspots. Nature. 2007;445:757–60.

13. IPBES. Global assessment report on biodiversity and ecosystem services of the Intergovernmental Science-Policy Platform on Biodiversity and Ecosystem Services. Bonn, Germany: IPBES secretariat; 2019.

14. Tucker CM, Aze T, Cadotte MW, Cantalapiedra JL, Chisholm C, Díaz S, et al. Assessing the utility of conserving evolutionary history. Biological Reviews. 2019;94(5):1740–60.

15. D’Costa C. Fifteen Footsteps Forward EDGE of Existence: Zoological Society of London; 2022 [Available from: https://www.edgeofexistence.org/blog/fifteen-footsteps-forward/.

16. ZSL. EDGE of Existence 2025 [Available from: https://www.edgeofexistence.org/.

17. Gumbs R, Gray CL, Böhm M, Burfield IJ, Couchman OR, Faith DP, et al. The EDGE2 protocol: Advancing the prioritisation of Evolutionarily Distinct and Globally Endangered species for practical conservation action. PLoS Biology. 2023;21(2):e3001991.

18. Faith DP. Conservation evaluation and phylogenetic diversity. Biological Conservation. 1992;61(1):1–10.

19. Redding DW, Mooers AO. Ranking Mammal Species for Conservation and the Loss of Both Phylogenetic and Trait Diversity. PloS ONE. 2015;10(12):e0141435.

20. Mazel F, Pennell MW, Cadotte MW, Diaz S, Dalla Riva GV, Grenyer R, et al. Prioritizing phylogenetic diversity captures functional diversity unreliably. Nature Communications. 2018;9(1):2888.

21. Mazel F, Mooers AO, Riva GVD, Pennell MW. Conserving Phylogenetic Diversity Can Be a Poor Strategy for Conserving Functional Diversity. Systematic Biology. 2017;66(6):1019–27.

22. Hidasi-Neto J, Loyola R, Cianciaruso MV. Global and local evolutionary and ecological distinctiveness of terrestrial mammals: identifying priorities across scales. Diversity and Distributions. 2015;21:548–59.

23. Cachera M, Le Loc’h F. Assessing the relationships between phylogenetic and functional singularities in sharks (Chondrichthyes). Ecology and Evolution. 2017;7(16):6292–303.

24. Devictor V, Mouillot D, Meynard C, Jiguet F, Thuiller W, Mouquet N. Spatial mismatch and congruence between taxonomic, phylogenetic and functional diversity: the need for integrative conservation strategies in a changing world. Ecology Letters. 2010;13(8):1030–40.

25. Dayan T, Simberloff D. Ecological and community-wide character displacement: the next generation. Ecology Letters. 2005;8(8):875–94.

26. Cadotte M, Albert CH, Walker SC. The ecology of differences: assessing community assembly with trait and evolutionary distances. Ecology Letters. 2013;16(10):1234–44.

27. Flynn DFB, Mirotchnick N, Jain M, Palmer MI, Naeem S. Functional and phylogenetic diversity as predictors of biodiversity–ecosystem-function relationships. Ecology. 2011;92(8):1573–81.

28. Pimiento C, Albouy C, Silvestro D, Mouton TL, Velez L, Mouillot D, et al. Functional diversity of sharks and rays is highly vulnerable and supported by unique species and locations worldwide. Nature Communications. 2023;14:7691.

29. Sánchez-Villagra MR, Horovitz I, Motokawa M. A comprehensive morphological analysis of talpid moles (Mammalia) phylogenetic relationships. Cladistics. 2006;22(1):59–88.

30. Linares-Martín A, Furió M, de Soler BG, Agustí J, Oms O, Grandi F, et al. An unexpected Scalopini mole (Talpidae, Mammalia) from the Pliocene of Europe sheds light on the phylogeny of talpids. Scientific Reports. 2025;15:Article number: 24928.

31. Dekker RWRJ. The distribution and status of nesting grounds of the Maleo Macrocephalon maleo in Sulawesi, Indonesia. Biological Conservation. 1990;51(2):139–50.

32. BirdLife International. Macrocephalon maleo 2021 [Available from: 10.2305/IUCN.UK.2021-3.RLTS.T22678576A194673255.en.

33. Indrawan M, Wahid N, Argeloo M, Mile-Doucet S, Tasirin J, Koh LP, et al. All politics is local: the case of Macrocephalon maleo conservation on Sulawesi, Indonesia. Brief Communication. 2012;21:3735–44.

34. Karim HA, Najib NN, Ayu SM, Fidel F. Characteristics of Maleo bird spawning nests (Macrocephalon maleo) in Lake Towuti, South Sulawesi, Indonesia. Biodiversitas Journal of Biological Diversity. 2023;24(2):690–6.

35. Hadly EA, Spaeth PA, Li C. Niche conservatism above the species level. Proceedings of the National Academy of Sciences. 2009;106:19707–14.

36. Olalla-Tárraga MÁ, McInnes L, Bini LM, Diniz-Filho JAF, Fritz SA, Hawkins BA, et al. Climatic niche conservatism and the evolutionary dynamics in species range boundaries: global congruence across mammals and amphibians. Journal of Biogeography. 2011;38(12):2237–47.

37. Long JA. Why Australasian vertebrate animals are so unique – A palaeontological perspective. General and Comparative Endocrinology. 2017;244:2–10.

38. García-Navas V, Westerman M. Niche conservatism and phylogenetic clustering in a tribe of arid-adapted marsupial mice, the Sminthopsini. Journal of Evolutionary Biology. 2018;31(8):1204–15.

39. Weir JT, Schluter D. Ice sheets promote speciation in boreal birds. Procedings of the Royal Society B. 2004;271(1551):1881–7.

40. Weir JT, Schluter D. The Latitudinal Gradient in Recent Speciation and Extinction Rates of Birds and Mammals. Science. 2007;315(5818):1574–6.

41. Pennell MW, FitzJohn RG, Cornwell WK, Harmon LJ. Model Adequacy and the Macroevolution of Angiosperm Functional Traits. The American Naturalist. 2015;186(2):E33–E50.

42. Tucker MA, Böhning-Gaese K, Fagan WF, Fryxell JM, Van Moorter B, Alberts SC, et al. Moving in the Anthropocene: Global reductions in terrestrial mammalian movements. Science. 2018;359:466–9.

43. Robuchon M, Silva Jd, Dubois G, Gumbs R, Hoban S, Laikre L, et al. Conserving species’ evolutionary potential and history: Opportunities under the Kunming–Montreal Global Biodiversity Framework. Conservation Science and Practice. 2023;5(6):e12929.

44. Pollock LJ, Thuiller W, Jetz W. Large conservation gains possible for global biodiversity facets. Nature. 2017;546(7656):141–4.

45. Owen NR, Gumbs R, Gray CL, Faith DP. Global conservation of phylogenetic diversity captures more than just functional diversity. Nature Communications. 2019;10:859.

46. Dee LE, Miller SJ, Peavey LE, Bradley D, Gentry RR, Startz R, et al. Functional diversity of catch mitigates negative effects of temperature variability on fisheries yields. Proceedings of the Royal Society B: Biological Sciences. 2016;283(1836):20161435.

47. Gagic V, Bartomeus I, Jonsson T, Taylor A, Winqvist C, Fischer C, et al. Functional identity and diversity of animals predict ecosystem functioning better than species-based indices. Proceedings of the Royal Society B: Biological Sciences. 2015;282(1801):20142620.

48. Lefcheck JS, Duffy JE. Multitrophic functional diversity predicts ecosystem functioning in experimental assemblages of estuarine consumers. Ecology. 2015;96(11):2973–83.

49. Huang X, Su J, Li S, Liu W, Lang X. Functional diversity drives ecosystem multifunctionality in a Pinus yunnanensis natural secondary forest. Scientific Reports. 2019;9(1):6979.

50. Taylor BW, Flecker AS, Hall RO. Loss of a harvested fish species disrupts carbon flow in a diverse tropical river. Science. 2006;313:833–6.

51. Faith DP. A general model for biodiversity and its value. In: Garson J, Plutynski A, Sarkar S, editors. The Routledge handbook of philosophy of biodiversity. New York: Routledge; 2017. p. 69-85.

52. Forest F, Moat J, Baloch E, Brummitt NA, Bachman SP, Ickert-Bond S, et al. Gymnosperms on the EDGE. Scientific Reports. 2018;8:6053.

53. Faith DP. Biodiversity and evolutionary history: useful extensions of the PD phylogenetic diversity assessment framework. Annals of the New York Academy of Sciences. 2013;1289:69–89.

54. Cooke RSC, Eigenbrod F, Bates AE. Ecological distinctiveness of birds and mammals at the global scale. Global Ecology and Conservation. 2020;22:e00970.

55. Faith DP. Valuation and Appreciation of Biodiversity: The “Maintenance of Options” Provided by the Variety of Life. Frontiers in Ecology and Evolution. 2021;9.

56. Chan KMA, Balvanera P, Benessaiah K, Chapman M, Díaz S, Gómez-Baggethun E, et al. Why protect nature? Rethinking values and the environment. Proceedings of the National Academy of Sciences. 2016;113(6):1462–5.

57. Marvier M, Kareiva P. Extinction is a moral wrong but conservation is complicated. Biological Conservation. 2014;176:281–2.

58. Marvier M, Wong H. Resurrecting the conservation movement. Journal of Environmental Studies and Sciences. 2012;2(4):291–5.

59. Pearson RG. Reasons to Conserve Nature. Trends in Ecology & Evolution. 2016;31(5):366–71.

60. Batavia C, Nelson MP. For goodness sake! What is intrinsic value and why should we care? Biological Conservation. 2017;209:366–76.

61. Piccolo JJ. Intrinsic values in nature: Objective good or simply half of an unhelpful dichotomy? Journal for Nature Conservation. 2017;37:8–11.

62. Jensen EL, Mooers AØ, Caccone A, Russello MA. I-HEDGE: determining the optimum complementary sets of taxa for conservation using evolutionary isolation. PeerJ. 2016;4:e2350.

63. Mouchet M, Guilhaumon F, Villéger S, Mason NWH, Tomasini J-A, Mouillot D. Towards a consensus for calculating dendrogram-based functional diversity indices. Oikos. 2008;117(5):794–800.

64. Petchey OL, O’Gorman EJ, Flynn DFB. A functional guide to functional diversity measures. In: Shahid N, Bunker DE, Hector A, Loreau M, Perrings C, editors. Biodiversity, Ecosystem Functioning, and Human Wellbeing: An Ecological and Economic Perspective. Oxford: Oxford Academic; 2009.

65. Kondratyeva A, Grandcolas P, Pavoine S. Reconciling the concepts and measures of diversity, rarity and originality in ecology and evolution. Biological Reviews. 2019;94(4):1317–37.

66. Pavoine S, Bonsall MB, Dupaix A, Jacob U, Ricotta C. From phylogenetic to functional originality: Guide through indices and new developments. Ecological Indicators. 2017;82:196–205.

67. Hidalgo-Cossio M, Salazar-Bravo J, Tarifa T. NUEVAS LOCALIDADES EN EL CENTRO DE BOLIVIA PARA LA ESPECIE ENDÉMICA Abrocoma boliviensis (RODENTIA: ABROCOMIDAE). Mastozoología Neotropical. 2016;23(1):165–70.

68. Nowak R. Walker’s Mammals of the World. London: Johns Hopkins Universtiy Press; 1999.

69. Opazo JC. A molecular timescale for caviomorph rodents (Mammalia, Hystricognathi). Molecular Phylogenetics and Evolution. 2005;37(3):932–7.

70. Tarifa T, Azurduy C, Vargas RR, Huanca N, Terán J, Arriaran G, et al. OBSERVATIONS ON THE NATURAL HISTORY OF Abrocoma sp. (RODENTIA, ABROCOMIDAE) IN A Polylepis WOODLAND IN BOLIVIA. Mastozoología Neotropical. 2009;16(1):253–8.

71. Froese GZL, Mustari AH. Assessments of Maleo Macrocephalon maleo nesting grounds in South-east Sulawesi reveal severely threatened populations. Bird Conservation International. 2019;29(4):497–502.

72. Steel M, Mimoto A, Mooers AØ. Hedging Our Bets: The Expected Contribution of Species to Future Phylogenetic Diversity. Evolutionary Bioinformatics. 2007;3.

73. Faith DP. Threatened species and the potential loss of phylogenetic diversity: Conservation scenarios based on estimated extinction probabilities and phylogenetic risk analysis. Conservation Biology. 2008;22(6):1461–70.

74. Faith DP, Reid CAM, Hunter J. Integrating phylogenetic diversity, complementarity, and endemism for conservation assessment. Conservation Biology. 2004;18(1):255–61.

75. Davis M, Faurby S, Svenning J-C. Mammal diversity will take millions of years to recover from the current biodiversity crisis. Proceedings of the National Academy of Sciences. 2018;115(44):11262–7.

76. Jones KE, Bielby J, Cardillo M, Fritz SA, O’Dell J, Orme CDL, et al. PanTHERIA: a species-level database of life history, ecology, and geography of extant and recently extinct mammals. Ecology. 2009;90:2648-.

77. Myhrvold NP, Baldridge E, Chan B, Sivam D, Freeman DL, Ernest SKM. An amniote life-history database to perform comparative analyses with birds, mammals, and reptiles. Ecology. 2015;96:3109-.

78. Wilman H, Belmaker J, Simpson J, de la Rosa C, Rivadeneira MM. EltonTraits 1.0: Species-level foraging attributes of the world’s birds and mammals. Ecology. 2014;95(7):2027.

79. R Core Team. R: A Language and Environment for Statistical Computing. Vienna, Austria: R Foundation for Statistical Computing; 2024.

80. Winemiller KO, Fitzgerald DB, Bower LM, Pianka ER. Functional traits, convergent evolution, and periodic tables of niches. Ecology Letters. 2015(18):737–51.

81. Violle C, Navas M-L, Vile D, Kazakou E, Fortunel C, Hummel I, et al. Let the concept of trait be functional! Oikos. 2007;116:882–92.

82. Cooke RSC, Bates AE, Eigenbrod F. Global trade-offs of functional redundancy and functional dispersion for birds and mammals. Global Ecology and Biogeography. 2019;28(4):484–95.

83. Humphries MM, McCann KS. Metabolic ecology. Journal of Animal Ecology. 2014;83(1):7–19.

84. Kembel SW, Cowan PD, Helmus MR, Cornwell WK, Morlon H, Ackerly DD, et al. Picante: R tools for integrating phylogenies and ecology. Bioinformatics. 2010;26:1463–4.

85. Upham NS, Esselstyn JA, Jetz W. Inferring the mammal tree: Species-level sets of phylogenies for questions in ecology, evolution, and conservation. PLoS Biology. 2019;12(12):e3000494.

86. Jetz W, Thomas GH, Joy JB, Hartmann K, Mooers AO. The global diversity of birds in space and time. Nature. 2012;491(7424):444–8.

87. Nakagawa S, De Villemereuil PA. General Method for Simultaneously Accounting for Phylogenetic and Species Sampling Uncertainty via Rubin’s Rules in Comparative Analysis. Systematic Biology. 2019;68(4):632–41.

88. Johnson TF, Isaac NJB, Paviolo A, González-Suárez M. Handling missing values in trait data. Global Ecology and Biogeography. 2021;30(1):51–62.

89. IUCN. IUCN Red List of Threatened Species. Version 2025-1 2025 [Available from: www.iucnredlist.org.

90. Orme D, Freckleton R, Thomas G, Petzoldt T, Fritz S, Isaac NJB, et al. caper: Comparative Analyses of Phylogenetics and Evolution in R. 2018.

91. Maechler M, Rousseeuw P, Struyf A, Hubert M, Hornik K. cluster: Cluster Analysis Basics and Extensions. 2022.

92. Maire E, Grenouillet G, Brosse S, Villéger S. How many dimensions are needed to accurately assess functional diversity? A pragmatic approach for assessing the quality of functional spaces. Global Ecology and Biogeography. 2015;24(6):728–40.

93. Bougeard S, Dray S. Supervised Multiblock Analysis in R with the ade4 Package. Journal of Statistical Software. 2018;86(1):1–17.

94. Chessel D, Dufour A, Thioulouse J. The ade4 Package – I: One-Table Methods. R News. 2004;4(1):5–10.

95. Dray S, Dufour A. The ade4 Package: Implementing the Duality Diagram for Ecologists. Journal of Statistical Software. 2007;20(4):1–20.

96. Dray S, Dufour A, Chessel D. The ade4 Package – II: Two-Table and K-Table Methods. R News. 2007;7(2):47–52.

97. Thioulouse J, Dray S, Dufour A, Siberchicot A, Jombart T, Pavoine S. Multivariate Analysis of Ecological Data with ade4: Springer; 2018.

98. Villéger S, Mason NWH, Mouillot D. New multidimensional functional diversity indices for a multifaceted framework in functional ecology. Ecology. 2008;89(8):2290–301.

99. Mouillot D, Graham NAJ, Villéger S, Mason NWH, Bellwood DR. A functional approach reveals community responses to disturbances. Trends in Ecology & Evolution. 2013;28(3):167–77.

100. Leitão RP, Zuanon J, Villéger S, Williams SE, Baraloto C, Fortunel C, et al. Rare species contribute disproportionately to the functional structure of species assemblages. Proceedings Biological Sciences. 2016;283(1828):20160084.

101. Griffin JN, Leprieur F, Silvestro D, Lefcheck JS, Albouy C, Rasher DB, et al. Functionally unique, specialised, and endangered (FUSE) species: Towards integrated metrics for the conservation prioritisation toolbox. bioRxiv. 2020.

102. Fritz SA, Purvis A. Selectivity in mammalian extinction risk and threat types: a new measure of phylogenetic signal strength in binary traits. Conservation Biology. 2010;24(4):1042–51.

103. 103. ZSL. Focal Species EDGE of Existencen.d. [Available from: https://www.edgeofexistence.org/focal-species/.

104. Pearse WD, Chase MW, Crawley MJ, Dolphin K, Fay MF, Joseph JA, et al. Beyond the EDGE with EDAM: Prioritising British Plant Species According to Evolutionary Distinctiveness, and Accuracy and Magnitude of Decline. PLoS ONE. 2015;10(5):e0126524.

105. Wilson HB, Joseph LN, Moore AL, Possingham HP. When should we save the most endangered species? Ecology Letters. 2011;14:886–90.

