## Supplemental Table 1 for "Integrating functional and phylogenetic perspectives reveals new priorities for bird and mammal conservation"

*Table S1* – Traits included in the functional dissimilarity matrices used to calculate FUn and FSp in FUSE and EFUSE. Trait data from the EltonTraits (3), PanTHERIA (4), Amniota (5)

| Trait | Description | Database |
|---|---|---|
| **Mammals** | | |
| **Body Mass** | Adult body mass (g) | EltonTraits |
| **Activity Cycle** | Activity cycle of each species measured for non-captive populations | PanTHERIA |
| **Trophic Level** | Trophic level of each species | PanTHERIA |
| **Litters Per Year** | Number of litters per female per year | PanTHERIA |
| **Habitat Breadth** | Number of habitat layers used | PanTHERIA |
| **Sexual Maturity Age** | Age when individuals are first physically capable of reproducing | PanTHERIA |
| **Basal Metabolic Rate** | Basal metabolic rate of adult individuals (mL.O2/hr) | PanTHERIA |
| **Diet Breadth** | Number of dietary categories eaten by each species | PanTHERIA |
| **Species Range** | Total extent of a species range (calculated using the total extent of a species range with a global equal-area projection (Mollweide)) (km2) | PanTHERIA |
| **Litter/clutch size** | The size of the litter/clutch | Amniota |
| **Birds** | | |
| **Habitat breadth** | Number of habitat layers used | Imputed |
| **Diet Category** | Assignment to the dominant among five diet categories based on the summed scores of constituent individual diets | EltonTraits |
| **Nocturnal** | Foraging activity at night | EltonTraits |
| **Body Mass** | Adult body mass (g) | EltonTraits |
| **Diet - Invertebrates** | Percent use of: Invertebrates | EltonTraits |
| **Diet - Vend** | Percent use of: Mammals, Birds | EltonTraits |
| **Diet - Vect** | Percent use of: Reptiles, snakes, amphibians, salamanders | EltonTraits |
| **Diet - Fish** | Percent use of: Fish | EltonTraits |
| **Diet - Vertebrates** | Percent use of: Vertebrates | EltonTraits |
| **Diet - Scavenge** | Percent use of: Scavenge, garbage, offal, carcasses, trawlers, carrrion | EltonTraits |
| **Diet - Fruit** | Percent use of: Fruit, drupes | EltonTraits |
| **Diet - Nectar** | Percent use of: Nectar, pollen, plant exudates, gums | EltonTraits |
| **Diet - Seeds** | Percent use of: Seed, maize, nuts, spores, wheat, grains | EltonTraits |
| **Diet - Plant (O)** | Percent use of: Other plant material, Grass, ground vegetation, seedlings, weeds, lichen, moss | EltonTraits |
| **Foraging below surface** | Prevalence of: Foraging below the water surfaces | EltonTraits |

| | | |
|---|---|---|
| **Foraging around surface** | Prevalence of: Foraging on or just (<5 inches) below water surface | EltonTraits |
| **Foraging on ground** | Prevalence of: Foraging on ground | EltonTraits |
| **Foraging in understory** | Prevalence of: Foraging below 2m in understory in forest, forest edges, bushes or shrubs | EltonTraits |
| **Foraging in mid - high levels in trees** | Prevalence of: Foraging in mid to high levels in trees or high bushes (2m upward), but below canopy | EltonTraits |
| **Foraging in canopy** | Prevalence of: Foraging in or just above (from) tree canopy | EltonTraits |
| **Aerial foraging** | Prevalence of: Foraging well above vegetation or any structures | EltonTraits |
| **Pelagic Specialist** | Pelagic foraging specialist | EltonTraits |
| **Litter/clutch size** | The size of the litter/clutch | Amniota |

4
