## Supplemental Table 2 for "Integrating functional and phylogenetic perspectives reveals new priorities for bird and mammal conservation"

*Table S2* – List of 100 mammals with the highest EFUSE scores (the EFUSE mammal top-100)

| Species | Common name | Order | EFUSE Score |
|---|---|---|---|
| ***Abrocoma boliviensis*** | Bolivian chinchilla rat | Rodentia | 4.90 |
| ***Abrocoma bennettii*** | Bennett's chinchilla rat | Rodentia | 3.54 |
| ***Bradypus pygmaeus*** | Pygmy three-toed sloth | Pilosa | 2.63 |
| ***Tokudaia muenninki*** | Muennink's spiny rat | Rodentia | 2.57 |
| ***Hypogeomys antimena*** | Malagasy giant rat | Rodentia | 2.54 |
| ***Desmana moschata*** | Russian desman | Soricomorpha | 2.51 |
| ***Zyzomys pedunculatus*** | Central rock rat | Rodentia | 2.48 |
| ***Neomonachus schauinslandi*** | Hawaiian monk seal | Carnivora | 2.47 |
| ***Macaca nigra*** | Celebes crested macaque | Primates | 2.37 |
| ***Addax nasomaculatus*** | Addax | Artiodactyla | 2.37 |
| ***Pongo abelii*** | Sumatran orangutan | Primates | 2.36 |
| ***Gorilla gorilla*** | Western gorilla | Primates | 2.36 |
| ***Gorilla beringei*** | Eastern gorilla | Primates | 2.35 |
| ***Dicerorhinus sumatrensis*** | Sumatran rhinoceros | Perissodactyla | 2.33 |
| ***Pongo pygmaeus*** | Bornean orangutan | Primates | 2.32 |
| ***Nanger dama*** | Dama gazelle | Artiodactyla | 2.30 |
| ***Macaca silenus*** | Lion-tailed macaque | Primates | 2.27 |
| ***Pteronura brasiliensis*** | Giant otter | Carnivora | 2.25 |
| ***Dorcopsis atrata*** | Black dorcopsis | Diprotodontia | 2.25 |
| ***Loxodonta africana*** | African bush elephant | Proboscidea | 2.23 |
| ***Manis culionensis*** | Philippine pangolin | Pholidota | 2.21 |
| ***Varecia rubra*** | Red ruffed lemur | Primates | 2.20 |
| ***Varecia variegata*** | Black-and-white ruffed lemur | Primates | 2.19 |
| ***Lycaon pictus*** | African wild dog | Carnivora | 2.16 |
| ***Cephalophus jentinki*** | Jentink's duiker | Artiodactyla | 2.15 |
| ***Rhinopithecus strykeri*** | Myanmar snub-nosed monkey | Primates | 2.14 |
| ***Lontra felina*** | Marine otter | Carnivora | 2.13 |
| ***Phocarctos hookeri*** | New Zealand sea lion | Carnivora | 2.13 |
| ***Zaglossus attenboroughi*** | Attenborough's long-beaked echidna | Monotremata | 2.12 |
| ***Mustela lutreola*** | European mink | Carnivora | 2.12 |
| ***Indri indri*** | Indri | Primates | 2.11 |
| ***Cebus kaapori*** | Kaapori capuchin | Primates | 2.10 |
| ***Urocitellus brunneus*** | Northern Idaho ground squirrel | Rodentia | 2.10 |
| ***Nasalis larvatus*** | Proboscis monkey | Primates | 2.10 |
| ***Hapalemur alaotrensis*** | Lac Alaotra bamboo lemur | Primates | 2.10 |
| ***Trachypithecus*** | Cat Ba langur | Primates | 2.10 |

| | | | |
|---|---|---|---|
| ***poliocephalus*** | | | |
| ***Diceros bicornis*** | Black rhinoceros | Perissodactyla | 2.10 |
| ***Mungotictis decemlineata*** | Narrow-striped mongoose | Carnivora | 2.09 |
| ***Marmota vancouverensis*** | Vancouver Island marmot | Rodentia | 2.08 |
| ***Cryptochloris wintoni*** | De Winton's golden mole | Afrosoricida | 2.08 |
| ***Rhinoceros sondaicus*** | Javan rhinoceros | Perissodactyla | 2.08 |
| ***Brachyteles arachnoides*** | Southern muriqui | Primates | 2.07 |
| ***Propithecus tattersalli*** | Golden-crowned sifaka | Primates | 2.07 |
| ***Brachyteles hypoxanthus*** | Northern muriqui | Primates | 2.06 |
| ***Sus cebifrons*** | Visayan warty pig | Artiodactyla | 2.05 |
| ***Macaca pagensis*** | Pagai Island macaque | Primates | 2.05 |
| ***Tympanoctomys loschalchalerosorum*** | Chalchalero viscacha rat | Rodentia | 2.04 |
| ***Manis javanica*** | Sunda pangolin | Pholidota | 2.04 |
| ***Choeropsis liberiensis*** | Pygmy hippopotamus | Artiodactyla | 2.04 |
| ***Manis pentadactyla*** | Chinese pangolin | Pholidota | 2.04 |
| ***Simias concolor*** | Pig-tailed langur | Primates | 2.04 |
| ***Rhinopithecus avunculus*** | Tonkin snub-nosed monkey | Primates | 2.02 |
| ***Bos sauveli*** | Kouprey | Artiodactyla | 2.01 |
| ***Peromyscus dickeyi*** | Dickey's deer mouse | Rodentia | 2.00 |
| ***Bradypus torquatus*** | Northern maned sloth | Pilosa | 2.00 |
| ***Lasiorhinus krefftii*** | Northern hairy-nosed wombat | Diprotodontia | 2.00 |
| ***Procyon pygmaeus*** | Cozumel raccoon | Carnivora | 2.00 |
| ***Sapajus xanthosternos*** | Golden-bellied capuchin | Primates | 2.00 |
| ***Tylomys tumbalensis*** | Tumbala climbing rat | Rodentia | 1.99 |
| ***Trachypithecus delacouri*** | Delacour's langur | Primates | 1.99 |
| ***Eubalaena glacialis*** | North Atlantic right whale | Cetacea | 1.98 |
| ***Propithecus coquereli*** | Coquerel's sifaka | Primates | 1.96 |
| ***Burramys parvus*** | Mountain pygmy possum | Diprotodontia | 1.96 |
| ***Bubalus mindorensis*** | Tamaraw | Artiodactyla | 1.96 |
| ***Abditomys latidens*** | Luzon broad-toothed rat | Rodentia | 1.95 |
| ***Propithecus perrieri*** | Perrier's sifaka | Primates | 1.95 |
| ***Equus africanus*** | African wild ass | Perissodactyla | 1.95 |
| ***Bos javanicus*** | Banteng | Artiodactyla | 1.94 |
| ***Zaglossus bruijnii*** | Western long-beaked echidna | Monotremata | 1.94 |
| ***Lontra provocax*** | Southern river otter | Carnivora | 1.93 |
| ***Geocapromys ingrahami*** | Bahamian hutia | Rodentia | 1.92 |
| ***Pan troglodytes*** | Chimpanzee | Primates | 1.92 |
| ***Beatragus hunteri*** | Hirola | Artiodactyla | 1.92 |
| ***Pan paniscus*** | Bonobo | Primates | 1.92 |
| ***Tylomys bullaris*** | Chiapan climbing rat | Rodentia | 1.91 |
| ***Cercocebus galeritus*** | Tana River mangabey | Primates | 1.90 |

| | | | |
|---|---|---|---|
| ***Moschus fuscus*** | Black musk deer | Artiodactyla | 1.90 |
| ***Dendrolagus scottae*** | Tenkile | Diprotodontia | 1.89 |
| ***Nomascus concolor*** | Black crested gibbon | Primates | 1.88 |
| ***Uromys imperator*** | Emperor rat | Rodentia | 1.87 |
| ***Lipotes vexillifer*** | Baiji | Cetacea | 1.87 |
| ***Acrobates pygmaeus*** | Feathertail glider | Diprotodontia | 1.87 |
| ***Zyzomys palatalis*** | Carpentarian rock rat | Rodentia | 1.86 |
| ***Elephas maximus*** | Asian elephant | Proboscidea | 1.86 |
| ***Prolemur simus*** | Greater bamboo lemur | Primates | 1.85 |
| ***Phocoena sinus*** | Vaquita | Cetacea | 1.85 |
| ***Pygathrix cinerea*** | Gray-shanked douc | Primates | 1.85 |
| ***Nomascus hainanus*** | Hainan black crested gibbon | Primates | 1.85 |
| ***Propithecus diadema*** | Diademed sifaka | Primates | 1.85 |
| ***Abeomelomys sevia*** | Highland brush mouse | Rodentia | 1.84 |
| ***Rhinopithecus brelichi*** | Gray snub-nosed monkey | Primates | 1.84 |
| ***Hapalemur aureus*** | Golden bamboo lemur | Primates | 1.83 |
| ***Camelus ferus*** | Wild Bactrian camel | Artiodactyla | 1.83 |
| ***Okapia johnstoni*** | Okapi | Artiodactyla | 1.83 |
| ***Muntiacus vuquangensis*** | Giant muntjac | Artiodactyla | 1.83 |
| ***Abrawayaomys ruschii*** | Ruschi's rat | Rodentia | 1.83 |
| ***Equus grevyi*** | Grévy's zebra | Perissodactyla | 1.82 |
| ***Pygathrix nigripes*** | Black-shanked douc | Primates | 1.82 |
| ***Pygathrix nemaeus*** | Red-shanked douc | Primates | 1.82 |
| ***Octodon pacificus*** | Pacific degu | Rodentia | 1.82 |

3
