## Supplementary Results & Discussion for "Integrating functional and phylogenetic perspectives reveals new priorities for bird and mammal conservation"

*EDGE and FUSE scores*

For mammals, EDGE scores ranged from a minimum of 1.36 (Black-clawed brush-furred rat, *Lophuromys melanonyx*, VU on the IUCN Red List), to a maximum of 5.01 (Bolivian chinchilla rat, *Abrocoma boliviensis*, CR on the IUCN Red List). The median EDGE score for mammals was 1.69. Species classified as NT or LC on the IUCN Red List are excluded from the EDGE framework and therefore do not receive EDGE scores. FUSE scores ranged from a minimum of 0 (3020 species all LC on the IUCN Red List) to a maximum of 0.989 (Russian desman, *Desmana moschata*, CR on the IUCN Red List). The median FUSE score for mammals was 0.

For birds, EDGE scores ranged from a minimum of 1.24 (Henderson reed warbler, *Acrocephalus taiti*, VU on the IUCN Red List), to a maximum of 4.47 (Beaudouin's snake eagle, *Circaetus beaudouini*, VU on the IUCN Red List). The median EDGE score for birds was 2.56. FUSE scores ranged from a minimum of 0 (6337 species all LC on the IUCN Red List) to a maximum of 0.781 (maleo, *Macrocephalon maleo*, CR on the IUCN Red List). The median FUSE score for birds was 0.

### Discussion

*EcoEDGE*

EcoEDGE is calculated as:

$$Score = ln[1 + (ED \times w_1 + EcoD \times w_2)] + \times\, ln(2)$$

where GE corresponds to the Red List category in which the species is categorised (LC=0, NT=1, VU=2, EN=3 and CR = 4). $w_1$ and $w_2$ are weights given to ED and EcoD, respectively. EcoD is ecological distinctiveness, calculated using the fair proportions method (1) for species in a functional dendrogram and ED is evolutionary distinctiveness, calculated using the fair proportions method for species in a given phylogeny (2).

### References

1. Redding, D.W. (2003) Incorporating genetic distinctness and reserve occupancy into a conservation prioritisation approach. Master's Thesis. University of East Anglia.

2. Hidasi-Neto, J., Loyola, R. & Cianciaruso, M.V. (2015) Global and local evolutionary and ecological distinctiveness of terrestrial mammals: identifying priorities across scales. Diversity and Distributions. 21, 548–559. doi:10.1111/ddi.12320.

3. Wilman, H., Belmaker, J., Simpson, J., de la Rosa, C., Rivadeneira, M.M. & Jetz, W. (2014) EltonTraits 1.0: Species-level foraging attributes of the world's birds and mammals. Ecology. 95 (7), 2027. doi:10.1890/13-1917.1.

4. Jones, K.E., Bielby, J., Cardillo, M., Fritz, S.A., O'Dell, J., et al. (2009) PanTHERIA: a species-level database of life history, ecology, and geography of extant and recently extinct mammals. Ecology. 90, 2648–2648. doi:10.1890/08-1494.1.

5. Myhrvold, N.P., Baldridge, E., Chan, B., Sivam, D., Freeman, D.L. & Ernest, S.K.M. (2015) An amniote life-history database to perform comparative analyses with birds, mammals, and reptiles. Ecology. 96, 3109–3109. doi:10.1890/15-0846R.1.
