## Supplemental Figure 1 for "Integrating functional and phylogenetic perspectives reveals new priorities for bird and mammal conservation"

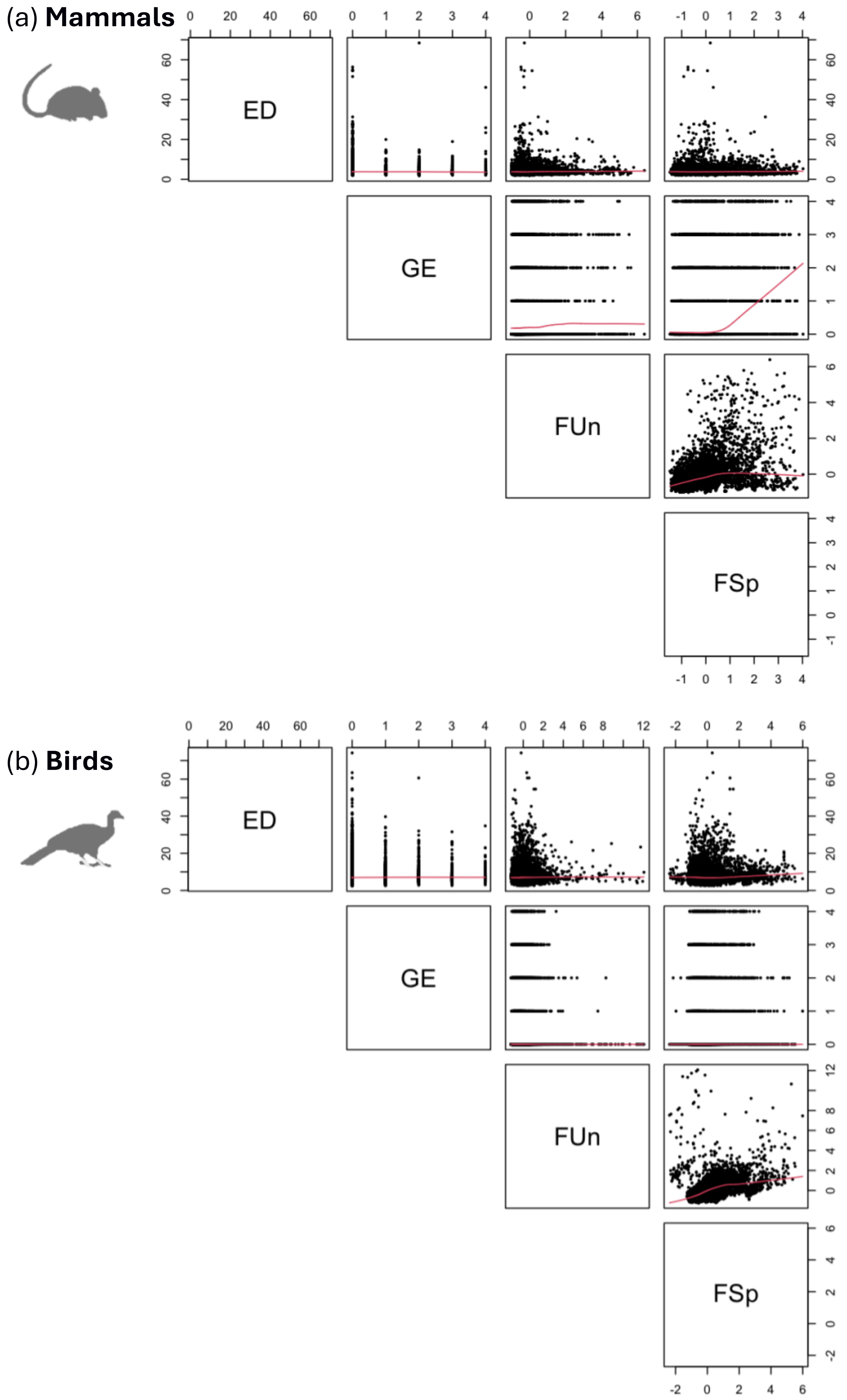


*Figure S1* – Scatter plots of the EDGE, FUSE, and EFUSE components against each other for (a) mammals and (b) birds. The Evolutionary Distinctiveness (ED) component of EDGE does not correlate strongly with either functional component (FUn and FSp) of FUSE, demonstrating that ED does not serve as a useful proxy for functional diversity.
