## Supplemental Table 7 for "Integrating functional and phylogenetic perspectives reveals new priorities for bird and mammal conservation"

*Table S7* – List of bird species which are on both the EDGE top-100 species and FUSE top-100 species. Rankings from highest score (1) to lowest.

| Species | Common name | EDGE ranking | FUSE ranking | EFUSE |
|---|---|---|---|---|
| ***Buteo ridgwayi*** | Ridgway's hawk | 14 | 89 | 7 |
| ***Anthracoceros montani*** | Sulu hornbill | 15 | 91 | 5 |
| ***Sagittarius serpentarius*** | Secretary bird | 18 | 90 | 45 |
| ***Spheniscus demersus*** | African penguin | 34 | 14 | 2 |
| ***Bubo shelleyi*** | Shelley's eagle-owl | 94 | 39 | 85 |
| ***Balearica regulorum*** | Grey crowned crane | 99 | 31 | 49 |
