## Supplemental Table 6 for "Integrating functional and phylogenetic perspectives reveals new priorities for bird and mammal conservation"

*Table S6* – List of 100 birds with the highest FUSE scores (the FUSE bird top-100)

| Species | Common name | Order | FUSE Score |
|---|---|---|---|
| ***Macrocephalon maleo*** | Maleo | Galliformes | 0.782 |
| ***Mergus octosetaceus*** | Brazilian merganser | Anseriformes | 0.680 |
| ***Otus insularis*** | Seychelles scops owl | Strigiformes | 0.674 |
| ***Ardeotis nigriceps*** | Great Indian bustard | Otidiformes | 0.652 |
| ***Gyps indicus*** | Indian vulture | Accipitriformes | 0.648 |
| ***Haliaeetus vociferoides*** | Madagascar fish eagle | Accipitriformes | 0.638 |
| ***Stagonopleura guttata*** | Diamond firetail | Passeriformes | 0.634 |
| ***Gymnogyps californianus*** | California condor | Accipitriformes | 0.627 |
| ***Necrosyrtes monachus*** | Hooded vulture | Accipitriformes | 0.621 |
| ***Glaucidium mooreorum*** | Pernambuco pygmy owl | Strigiformes | 0.603 |
| ***Pithecophaga jefferyi*** | Philippine eagle | Accipitriformes | 0.602 |
| ***Anas laysanensis*** | Laysan duck | Anseriformes | 0.576 |
| ***Aythya innotata*** | Madagascar pochard | Anseriformes | 0.569 |
| ***Spheniscus demersus*** | African penguin | Sphenisciformes | 0.568 |
| ***Diomedea dabbenena*** | Tristan albatross | Procellariiformes | 0.562 |
| ***Apteryx australis*** | Southern brown kiwi | Apterygiformes | 0.557 |
| ***Aegotheles savesi*** | New Caledonian owlet-nightjar | Apodiformes | 0.557 |
| ***Aythya baeri*** | Baer's pochard | Anseriformes | 0.552 |
| ***Aepypodius bruijnii*** | Waigeo brushturkey | Galliformes | 0.543 |
| ***Pauxi unicornis*** | Horned curassow | Galliformes | 0.538 |
| ***Rhinoplax vigil*** | Helmeted hornbill | Bucerotiformes | 0.537 |
| ***Gyps tenuirostris*** | Slender-billed vulture | Accipitriformes | 0.536 |
| ***Houbaropsis bengalensis*** | Bengal florican | Otidiformes | 0.534 |
| ***Mergus squamatus*** | Scaly-sided merganser | Anseriformes | 0.534 |
| ***Otus siaoensis*** | Siau scops owl | Strigiformes | 0.528 |
| ***Ardea insignis*** | White-bellied heron | Pelecaniformes | 0.527 |
| ***Megadyptes antipodes*** | Yellow-eyed penguin | Sphenisciformes | 0.527 |
| ***Gyps africanus*** | White-backed vulture | Accipitriformes | 0.525 |
| ***Sarcogyps calvus*** | Red-headed vulture | Accipitriformes | 0.521 |
| ***Thaumatibis gigantea*** | Giant ibis | Pelecaniformes | 0.517 |
| ***Balearica regulorum*** | Grey crowned crane | Gruiformes | 0.513 |
| ***Trigonoceps occipitalis*** | White-headed vulture | Accipitriformes | 0.511 |
| ***Gyps bengalensis*** | White-rumped vulture | Accipitriformes | 0.509 |
| ***Otis tarda*** | Great bustard | Otidiformes | 0.507 |
| ***Bubo philippensis*** | Philippine eagle-owl | Strigiformes | 0.507 |
| ***Crax alberti*** | Blue-billed curassow | Galliformes | 0.506 |
| ***Neotis ludwigii*** | Ludwig's bustard | Otidiformes | 0.505 |
| ***Leipoa ocellata*** | Malleefowl | Galliformes | 0.504 |

| | | | |
|---|---|---|---|
| ***Bubo shelleyi*** | Shelley's eagle-owl | Strigiformes | 0.501 |
| ***Heliopais personatus*** | Masked finfoot | Gruiformes | 0.500 |
| ***Grus americana*** | Whooping crane | Gruiformes | 0.499 |
| ***Apteryx mantelli*** | North Island brown kiwi | Apterygiformes | 0.497 |
| ***Terathopius ecaudatus*** | Bateleur | Accipitriformes | 0.492 |
| ***Phoebastria irrorata*** | Waved albatross | Procellariiformes | 0.490 |
| ***Apteryx haastii*** | Great spotted kiwi | Apterygiformes | 0.489 |
| ***Diomedea antipodensis*** | Antipodean albatross | Procellariiformes | 0.485 |
| ***Pipile pipile*** | Trinidad piping guan | Galliformes | 0.480 |
| ***Morus capensis*** | Cape gannet | Suliformes | 0.479 |
| ***Scotopelia ussheri*** | Rufous fishing owl | Strigiformes | 0.476 |
| ***Phalacrocorax capensis*** | Cape cormorant | Suliformes | 0.474 |
| ***Polemaetus bellicosus*** | Martial eagle | Accipitriformes | 0.474 |
| ***Thinornis novaeseelandiae*** | Shore plover | Charadriiformes | 0.471 |
| ***Pterodroma phaeopygia*** | Galápagos petrel | Procellariiformes | 0.465 |
| ***Diomedea amsterdamensis*** | Amsterdam albatross | Procellariiformes | 0.464 |
| ***Eudyptes moseleyi*** | Northern rockhopper penguin | Sphenisciformes | 0.457 |
| ***Pseudibis davisoni*** | White-shouldered ibis | Pelecaniformes | 0.454 |
| ***Eudyptes sclateri*** | Erect-crested penguin | Sphenisciformes | 0.452 |
| ***Didunculus strigirostris*** | Tooth-billed pigeon | Columbiformes | 0.452 |
| ***Otus thilohoffmanni*** | Serendib scops owl | Strigiformes | 0.451 |
| ***Pavo muticus*** | Green peafowl | Galliformes | 0.450 |
| ***Haliaeetus leucoryphus*** | Pallas's fish eagle | Accipitriformes | 0.447 |
| ***Porphyrio hochstetteri*** | Takahe | Gruiformes | 0.447 |
| ***Emberiza aureola*** | Yellow-breasted bunting | Passeriformes | 0.446 |
| ***Amazona auropalliata*** | Yellow-naped amazon | Psittaciformes | 0.446 |
| ***Spizaetus isidori*** | Black-and-chestnut eagle | Accipitriformes | 0.445 |
| ***Otus capnodes*** | Anjouan scops owl | Strigiformes | 0.444 |
| ***Pterodroma sandwichensis*** | Hawaiian petrel | Procellariiformes | 0.443 |
| ***Alcedo euryzona*** | Javan blue-banded kingfisher | Coraciiformes | 0.441 |
| ***Lophura edwardsi*** | Edwards's pheasant | Galliformes | 0.441 |
| ***Otus alfredi*** | Flores scops owl | Strigiformes | 0.441 |
| ***Lophura erythrophthalma*** | Crestless fireback | Galliformes | 0.438 |
| ***Torgos tracheliotos*** | Lappet-faced vulture | Accipitriformes | 0.437 |
| ***Aquila nipalensis*** | Steppe eagle | Accipitriformes | 0.434 |
| ***Ciconia stormi*** | Storm's stork | Ciconiiformes | 0.433 |
| ***Ciconia boyciana*** | Oriental stork | Ciconiiformes | 0.431 |
| ***Neophron percnopterus*** | Egyptian vulture | Accipitriformes | 0.431 |

| | | | |
|---|---|---|---|
| ***Diomedea sanfordi*** | Northern royal albatross | Procellariiformes | 0.431 |
| ***Cacatua sulphurea*** | Yellow-crested cockatoo | Psittaciformes | 0.430 |
| ***Ninox sumbaensis*** | Least boobook | Strigiformes | 0.430 |
| ***Ara ambiguus*** | Great green macaw | Psittaciformes | 0.430 |
| ***Otus pauliani*** | Karthala scops owl | Strigiformes | 0.428 |
| ***Thalassarche chrysostoma*** | Grey-headed albatross | Procellariiformes | 0.428 |
| ***Spheniscus mendiculus*** | Galapagos penguin | Sphenisciformes | 0.428 |
| ***Centrocercus minimus*** | Gunnison sage-grouse | Galliformes | 0.426 |
| ***Ara rubrogenys*** | Red-fronted macaw | Psittaciformes | 0.425 |
| ***Lathamus discolor*** | Swift parrot | Psittaciformes | 0.424 |
| ***Oxyura leucocephala*** | White-headed duck | Anseriformes | 0.421 |
| ***Tyto soumagnei*** | Red owl | Strigiformes | 0.420 |
| ***Buteo ridgwayi*** | Ridgway's hawk | Accipitriformes | 0.419 |
| ***Sagittarius serpentarius*** | Secretary bird | Accipitriformes | 0.418 |
| ***Anthracoceros montani*** | Sulu hornbill | Bucerotiformes | 0.418 |
| ***Otus ireneae*** | Sokoke scops owl | Strigiformes | 0.416 |
| ***Vultur gryphus*** | Andean condor | Accipitriformes | 0.415 |
| ***Falco cherrug*** | Saker falcon | Falconiformes | 0.413 |
| ***Puffinus newelli*** | Newell's shearwater | Procellariiformes | 0.412 |
| ***Circus maurus*** | Black harrier | Accipitriformes | 0.412 |
| ***Eutriorchis astur*** | Madagascar serpent eagle | Accipitriformes | 0.409 |
| ***Rallus wetmorei*** | Plain-flanked rail | Gruiformes | 0.407 |
| ***Papasula abbotti*** | Abbott's booby | Suliformes | 0.407 |
| ***Otus moheliensis*** | Moheli scops owl | Strigiformes | 0.406 |

3
