## Supplemental Table 5 for "Integrating functional and phylogenetic perspectives reveals new priorities for bird and mammal conservation"

*Table S5* – List of 100 birds with the highest EFUSE scores (the EFUSE bird top-100)

| Species | Common name | Order | EFUSE Score |
|---|---|---|---|
| ***Macrocephalon maleo*** | Maleo | Galliformes | 3.48 |
| ***Spheniscus demersus*** | African penguin | Sphenisciformes | 3.32 |
| ***Rowettia goughensis*** | Gough finch | Passeriformes | 3.27 |
| ***Otus insularis*** | Seychelles scops owl | Strigiformes | 3.25 |
| ***Anthracoceros montani*** | Sulu hornbill | Bucerotiformes | 3.22 |
| ***Mergus octosetaceus*** | Brazilian merganser | Anseriformes | 3.20 |
| ***Buteo ridgwayi*** | Ridgway's hawk | Accipitriformes | 3.19 |
| ***Gyps indicus*** | Indian vulture | Accipitriformes | 3.18 |
| ***Ardeotis nigriceps*** | Great Indian bustard | Otidiformes | 3.18 |
| ***Haliaeetus vociferoides*** | Madagascar fish eagle | Accipitriformes | 3.17 |
| ***Aythya innotata*** | Madagascar pochard | Anseriformes | 3.16 |
| ***Gymnogyps californianus*** | California condor | Accipitriformes | 3.15 |
| ***Glaucidium mooreorum*** | Pernambuco pygmy owl | Strigiformes | 3.12 |
| ***Aythya baeri*** | Baer's pochard | Anseriformes | 3.11 |
| ***Necrosyrtes monachus*** | Hooded vulture | Accipitriformes | 3.08 |
| ***Circaetusbeaudouini*** | Beaudouin's snake eagle | Accipitriformes | 3.05 |
| ***Pithecophaga jefferyi*** | Philippine eagle | Accipitriformes | 2.99 |
| ***Anas laysanensis*** | Anas laysanensis | Anseriformes | 2.98 |
| ***Calyptura cristata*** | Kinglet calyptura | Passeriformes | 2.97 |
| ***Diomedea dabbenena*** | Tristan albatross | Procellariiformes | 2.96 |
| ***Cacatua sulphurea*** | Yellow-crested cockatoo | Psittaciformes | 2.96 |
| ***Gyps tenuirostris*** | Slender-billed vulture | Accipitriformes | 2.95 |
| ***Gyps africanus*** | White-backed vulture | Accipitriformes | 2.95 |
| ***Houbaropsis bengalensis*** | Bengal florican | Otidiformes | 2.93 |
| ***Thaumatibis gigantea*** | Giant ibis | Pelecaniformes | 2.92 |
| ***Heliopais personatus*** | Masked finfoot | Gruiformes | 2.91 |
| ***Chondrohierax wilsonii*** | Cuban kite | Accipitriformes | 2.91 |
| ***Stagonopleura guttata*** | Diamond firetail | Passeriformes | 2.91 |
| ***Rhinoplax vigil*** | Helmeted hornbill | Bucerotiformes | 2.91 |
| ***Campephilus imperialis*** | Imperial woodpecker | Piciformes | 2.90 |
| ***Gyps bengalensis*** | White-rumped vulture | Accipitriformes | 2.90 |
| ***Sarcogyps calvus*** | Red-headed vulture | Accipitriformes | 2.89 |
| ***Pauxi unicornis*** | Horned curassow | Galliformes | 2.88 |
| ***Bostrychia bocagei*** | São Tomé ibis | Pelecaniformes | 2.88 |
| ***Campephilus principalis*** | Ivory-billed woodpecker | Piciformes | 2.88 |
| ***Otus siaoensis*** | Siau scops owl | Strigiformes | 2.88 |
| ***Ardea insignis*** | White-bellied heron | Pelecaniformes | 2.87 |
| ***Cacatua haematuropygia*** | Red-vented cockatoo | Psittaciformes | 2.87 |
| ***Cinclodes palliatus*** | White-bellied cinclodes | Passeriformes | 2.85 |
| ***Trigonoceps occipitalis*** | White-headed vulture | Accipitriformes | 2.84 |

| | | | |
|---|---|---|---|
| ***Crax alberti*** | Blue-billed curassow | Galliformes | 2.84 |
| ***Aegotheles savesi*** | New Caledonian owlet-nightjar | Apodiformes | 2.83 |
| ***Ara rubrogenys*** | Red-fronted macaw | Psittaciformes | 2.81 |
| ***Phoebastria irrorata*** | Waved albatross | Procellariiformes | 2.80 |
| ***Sagittarius serpentarius*** | Secretarybird | Accipitriformes | 2.78 |
| ***Lophura edwardsi*** | Edwards's pheasant | Galliformes | 2.77 |
| ***Lophura erythrophthalma*** | Crestless fireback | Galliformes | 2.76 |
| ***Lathamus discolor*** | Swift parrot | Psittaciformes | 2.75 |
| ***Balearica regulorum*** | Grey crowned crane | Gruiformes | 2.75 |
| ***Ara ambiguus*** | Great green macaw | Psittaciformes | 2.75 |
| ***Amazona auropalliata*** | Yellow-naped amazon | Psittaciformes | 2.75 |
| ***Pterodroma phaeopygia*** | Galápagos petrel | Procellariiformes | 2.73 |
| ***Campylopterus phainopeplus*** | Santa Marta sabrewing | Apodiformes | 2.73 |
| ***Didunculus strigirostris*** | Tooth-billed pigeon | Columbiformes | 2.72 |
| ***Cyanoramphus malherbi*** | Orange-fronted parakeet | Psittaciformes | 2.72 |
| ***Pipile pipile*** | Trinidad piping guan | Galliformes | 2.72 |
| ***Alcedo euryzona*** | Javan blue-banded kingfisher | Coraciiformes | 2.71 |
| ***Pseudibis davisoni*** | White-shouldered ibis | Pelecaniformes | 2.71 |
| ***Sypheotides indicus*** | Lesser florican | Otidiformes | 2.70 |
| ***Amazona imperialis*** | Imperial amazon | Psittaciformes | 2.70 |
| ***Emberiza aureola*** | Yellow-breasted bunting | Passeriformes | 2.70 |
| ***Numenius tenuirostris*** | Slender-billed curlew | Charadriiformes | 2.70 |
| ***Callaeas cinereus*** | South Island kōkako | Passeriformes | 2.69 |
| ***Vini ultramarina*** | Ultramarine lorikeet | Psittaciformes | 2.69 |
| ***Corvus kubaryi*** | Mariana crow | Passeriformes | 2.68 |
| ***Palmeria dolei*** | ʻAkohekohe | Passeriformes | 2.68 |
| ***Pezoporus occidentalis*** | Night parrot | Psittaciformes | 2.66 |
| ***Leptotila wellsi*** | Grenada dove | Columbiformes | 2.66 |
| ***Telespiza ultima*** | Nīhoa finch | Passeriformes | 2.66 |
| ***Ara glaucogularis*** | Blue-throated macaw | Psittaciformes | 2.65 |
| ***Cyanolimnas cerverai*** | Zapata rail | Gruiformes | 2.65 |
| ***Neophema chrysogaster*** | Orange-bellied parrot | Psittaciformes | 2.65 |
| ***Zosterops rotensis*** | Rota white-eye | Passeriformes | 2.65 |
| ***Rynchops albicollis*** | Indian skimmer | Charadriiformes | 2.65 |
| ***Cacatua alba*** | White cockatoo | Psittaciformes | 2.64 |
| ***Puffinus newelli*** | Newell's shearwater | Procellariiformes | 2.64 |
| ***Accipiter gundlachi*** | Gundlach's hawk | Accipitriformes | 2.64 |
| ***Carpococcyx viridis*** | Sumatran ground cuckoo | Cuculiformes | 2.63 |
| ***Terenura sicki*** | Orange-bellied antwren | Passeriformes | 2.63 |
| ***Charmosyna amabilis*** | Red-throated lorikeet | Psittaciformes | 2.63 |
| ***Podiceps gallardoi*** | Hooded grebe | Podicipediformes | 2.61 |

| | | | |
|---|---|---|---|
| ***Corvus unicolor*** | Banggai crow | Passeriformes | 2.61 |
| ***Pterodroma caribbaea*** | Jamaican petrel | Procellariiformes | 2.61 |
| ***Charadrius obscurus*** | New Zealand dotterel | Charadriiformes | 2.59 |
| ***Bubo shelleyi*** | Shelley's eagle-owl | Strigiformes | 2.59 |
| ***Puffinus auricularis*** | Townsend's shearwater | Procellariiformes | 2.59 |
| ***Zosterops nehrkorni*** | Sangihe white-eye | Passeriformes | 2.59 |
| ***Rostratula australis*** | Australian painted-snipe | Charadriiformes | 2.58 |
| ***Himantopus novaezelandiae*** | Black stilt | Charadriiformes | 2.58 |
| ***Mimus graysoni*** | Socorro mockingbird | Passeriformes | 2.58 |
| ***Numenius borealis*** | Eskimo curlew | Charadriiformes | 2.58 |
| ***Puffinus mauretanicus*** | Balearic shearwater | Procellariiformes | 2.57 |
| ***Cissa thalassina*** | Javan green magpie | Passeriformes | 2.57 |
| ***Columba argentina*** | Silvery pigeon | Columbiformes | 2.56 |
| ***Lophornis brachylophus*** | Short-crested coquette | Apodiformes | 2.56 |
| ***Gymnomyza aubryana*** | Crow honeyeater | Passeriformes | 2.55 |
| ***Charmosyna diadema*** | New Caledonian lorikeet | Psittaciformes | 2.55 |
| ***Vermivora bachmanii*** | Bachman's warbler | Passeriformes | 2.55 |
| ***Loxioides bailleui*** | Palila | Passeriformes | 2.54 |
| ***Bubo philippensis*** | Philippine eagle-owl | Strigiformes | 2.54 |

3
