## Supplemental Table 4 for "Integrating functional and phylogenetic perspectives reveals new priorities for bird and mammal conservation"

*Table S4* – List of mammal species which are on both the EDGE top-100 species and FUSE top-100 species. Rankings from highest score (1) to lowest.

| Species | Common name | EDGE ranking | FUSE ranking | EFUSE ranking |
|---|---|---|---|---|
| ***Lycaon pictus*** | African wild dog | 47 | 39 | 24 |
| ***Nasalis larvatus*** | Proboscis monkey | 48 | 51 | 34 |
| ***Addax nasomaculatus*** | Addax | 49 | 46 | 10 |
| ***Hypogeomys antimena*** | Malagasy giant rat | 55 | 11 | 5 |
| ***Moschus fuscus*** | Black musk deer | 69 | 89 | 77 |
| ***Neomonachus schauinslandi*** | Hawaiian monk seal | 89 | 4 | 8 |
| ***Cephalophus jentinki*** | Jentink's duiker | 96 | 27 | 25 |
