## Supplemental Table 3 for "Integrating functional and phylogenetic perspectives reveals new priorities for bird and mammal conservation"

*Table S3* – List of 100 mammals with the highest FUSE scores (the FUSE mammal top-100)

| Species | Common name | Order | FUSE Score |
|---|---|---|---|
| ***Desmana moschata*** | Russian desman | Soricomorpha | 0.989 |
| ***Pongo abelii*** | Sumatran orangutan | Primates | 0.982 |
| ***Pteronura brasiliensis*** | Giant otter | Carnivora | 0.963 |
| ***Neomonachus schauinslandi*** | Hawaiian monk seal | Carnivora | 0.955 |
| ***Pongo pygmaeus*** | Bornean orangutan | Primates | 0.946 |
| ***Macaca silenus*** | Lion-tailed macaque | Primates | 0.944 |
| ***Loxodonta africana*** | African bush elephant | Proboscidea | 0.936 |
| ***Dicerorhinus sumatrensis*** | Sumatran rhinoceros | Perissodactyla | 0.920 |
| ***Phocarctos hookeri*** | New Zealand sea lion | Carnivora | 0.917 |
| ***Macaca nigra*** | Celebes crested macaque | Primates | 0.908 |
| ***Hypogeomys antimena*** | Malagasy giant rat | Rodentia | 0.898 |
| ***Choeropsis liberiensis*** | Pygmy hippopotamus | Artiodactyla | 0.875 |
| ***Gorilla beringei*** | Eastern gorilla | Primates | 0.849 |
| ***Lontra felina*** | Marine otter | Carnivora | 0.844 |
| ***Nanger dama*** | Dama gazelle | Artiodactyla | 0.841 |
| ***Gorilla gorilla*** | Western gorilla | Primates | 0.835 |
| ***Manis culionensis*** | Philippine pangolin | Pholidota | 0.830 |
| ***Rhinopithecus strykeri*** | Myanmar snub-nosed monkey | Primates | 0.830 |
| ***Mungotictis decemlineata*** | Narrow-striped mongoose | Carnivora | 0.820 |
| ***Pan troglodytes*** | Chimpanzee | Primates | 0.819 |
| ***Pan paniscus*** | Bonobo | Primates | 0.817 |
| ***Lontra provocax*** | Southern river otter | Carnivora | 0.817 |
| ***Varecia rubra*** | Red ruffed lemur | Primates | 0.810 |
| ***Bradypus torquatus*** | Northern maned sloth | Pilosa | 0.797 |
| ***Hapalemur alaotrensis*** | Lac Alaotra bamboo lemur | Primates | 0.793 |
| ***Varecia variegata*** | Black-and-white ruffed lemur | Primates | 0.783 |
| ***Cephalophus jentinki*** | Jentink's duiker | Artiodactyla | 0.775 |
| ***Dorcopsis atrata*** | Black dorcopsis | Diprotodontia | 0.773 |
| ***Propithecus tattersalli*** | Golden-crowned sifaka | Primates | 0.771 |
| ***Pusa caspica*** | Caspian seal | Carnivora | 0.764 |
| ***Urocitellus brunneus*** | Northern Idaho ground squirrel | Rodentia | 0.762 |
| ***Rhinoceros sondaicus*** | Javan rhinoceros | Perissodactyla | 0.760 |
| ***Indri indri*** | Indri | Primates | 0.752 |
| ***Diceros bicornis*** | Black rhinoceros | Perissodactyla | 0.751 |
| ***Simias concolor*** | Pig-tailed langur | Primates | 0.75 |
| ***Elephas maximus*** | Asian elephant | Proboscidea | 0.743 |

| | | | |
|---|---|---|---|
| ***Marmota vancouverensis*** | Vancouver Island marmot | Rodentia | 0.74 |
| ***Mustela lutreola*** | European mink | Carnivora | 0.739 |
| ***Lycaon pictus*** | African wild dog | Carnivora | 0.739 |
| ***Sus cebifrons*** | Visayan warty pig | Artiodactyla | 0.733 |
| ***Bradypus pygmaeus*** | Pygmy three-toed sloth | Pilosa | 0.731 |
| ***Rhinopithecus avunculus*** | Tonkin snub-nosed monkey | Primates | 0.721 |
| ***Manis javanica*** | Sunda pangolin | Pholidota | 0.721 |
| ***Manis pentadactyla*** | Chinese pangolin | Pholidota | 0.721 |
| ***Canis simensis*** | Ethiopian wolf | Carnivora | 0.720 |
| ***Addax nasomaculatus*** | Addax | Artiodactyla | 0.719 |
| ***Equus grevyi*** | Grévy's zebra | Perissodactyla | 0.718 |
| ***Rhinoceros unicornis*** | Indian rhinoceros | Perissodactyla | 0.712 |
| ***Sus verrucosus*** | Javan warty pig | Artiodactyla | 0.712 |
| ***Sapajus xanthosternos*** | Golden-bellied capuchin | Primates | 0.711 |
| ***Nasalis larvatus*** | Proboscis monkey | Primates | 0.704 |
| ***Zaglossus attenboroughi*** | Attenborough's long-beaked echidna | Monotremata | 0.701 |
| ***Propithecus coquereli*** | Coquerel's sifaka | Primates | 0.701 |
| ***Panthera tigris*** | Tiger | Carnivora | 0.701 |
| ***Okapia johnstoni*** | Okapi | Artiodactyla | 0.694 |
| ***Enhydra lutris*** | Sea otter | Carnivora | 0.694 |
| ***Lasiorhinus krefftii*** | Northern hairy-nosed wombat | Diprotodontia | 0.688 |
| ***Peromyscus dickeyi*** | Dickey's deer mouse | Rodentia | 0.681 |
| ***Ursus maritimus*** | Polar bear | Carnivora | 0.681 |
| ***Brachyteles hypoxanthus*** | Northern muriqui | Primates | 0.680 |
| ***Procyon pygmaeus*** | Cozumel raccoon | Carnivora | 0.677 |
| ***Brachyteles arachnoides*** | Southern muriqui | Primates | 0.677 |
| ***Macaca pagensis*** | Pagai Island macaque | Primates | 0.670 |
| ***Cryptochloris wintoni*** | De Winton's golden mole | Afrosoricida | 0.666 |
| ***Rhinopithecus bieti*** | Black-and-white snub-nosed monkey | Primates | 0.666 |
| ***Propithecus perrieri*** | Perrier's sifaka | Primates | 0.663 |
| ***Eubalaena glacialis*** | North Atlantic right whale | Cetacea | 0.661 |
| ***Cebus kaapori*** | Kaapori capuchin | Primates | 0.650 |
| ***Chinchilla chinchilla*** | Short-tailed chinchilla | Rodentia | 0.638 |
| ***Catagonus wagneri*** | Chacoan peccary | Artiodactyla | 0.638 |
| ***Trachypithecus poliocephalus*** | Cat Ba langur | Primates | 0.634 |
| ***Geomys tropicalis*** | Tropical pocket gopher | Rodentia | 0.630 |
| ***Cephalophus spadix*** | Abbott's duiker | Artiodactyla | 0.629 |
| ***Equus africanus*** | African wild ass | Perissodactyla | 0.627 |
| ***Cercocebus galeritus*** | Tana River mangabey | Primates | 0.625 |
| ***Dendrolagus scottae*** | Tenkile | Diprotodontia | 0.622 |

| | | | |
|---|---|---|---|
| ***Chinchilla lanigera*** | Long-tailed chinchilla | Rodentia | 0.621 |
| ***Hippopotamus amphibius*** | Hippopotamus | Artiodactyla | 0.619 |
| ***Prionailurus planiceps*** | Flat-headed cat | Carnivora | 0.615 |
| ***Lipotes vexillifer*** | Baiji | Cetacea | 0.613 |
| ***Pygathrix cinerea*** | Gray-shanked douc | Primates | 0.605 |
| ***Bubalus depressicornis*** | Lowland anoa | Artiodactyla | 0.604 |
| ***Cynomys mexicanus*** | Mexican prairie dog | Rodentia | 0.602 |
| ***Hylobates klossii*** | Kloss's gibbon | Primates | 0.601 |
| ***Myrmecophaga tridactyla*** | Giant anteater | Pilosa | 0.600 |
| ***Rhinopithecus brelichi*** | Gray snub-nosed monkey | Primates | 0.600 |
| ***Bos sauveli*** | Kouprey | Artiodactyla | 0.597 |
| ***Bubalus mindorensis*** | Tamaraw | Artiodactyla | 0.596 |
| ***Moschus fuscus*** | Black musk deer | Artiodactyla | 0.594 |
| ***Propithecus diadema*** | Diademed sifaka | Primates | 0.592 |
| ***Porcula salvania*** | Pygmy hog | Artiodactyla | 0.591 |
| ***Burramys parvus*** | Mountain pygmy possum | Diprotodontia | 0.589 |
| ***Bubalus quarlesi*** | Mountain anoa | Artiodactyla | 0.589 |
| ***Manis crassicaudata*** | Indian pangolin | Pholidota | 0.589 |
| ***Trachypithecus delacouri*** | Delacour's langur | Primates | 0.584 |
| ***Prolemur simus*** | Greater bamboo lemur | Primates | 0.584 |
| ***Pygathrix nigripes*** | Black-shanked douc | Primates | 0.577 |
| ***Dugong dugon*** | Dugong | Sirenia | 0.577 |
| ***Tapirus indicus*** | Malayan tapir | Perissodactyla | 0.576 |
| ***Pygathrix nemaeus*** | Red-shanked douc | Primates | 0.576 |

3
